# Decentralizing cell-free RNA sensing with the use of low-cost cell extracts

**DOI:** 10.1101/2021.05.29.446205

**Authors:** Anibal Arce, Fernando Guzman Chavez, Chiara Gandini, Juan Puig, Tamara Matute, Jim Haseloff, Neil Dalchau, Jenny Molloy, Keith Pardee, Fernán Federici

**Author notes:** **Correspondence: Fernán Federici**.

## Abstract

Cell-free gene expression systems have emerged as a promising platform for field-deployed biosensing and diagnostics. When combined with programmable toehold switch-based RNA sensors, these systems can be used to detect arbitrary RNAs and freeze-dried for room temperature transport to the point-of-need. These sensors, however, have been implemented using reconstituted PURE cell-free protein expression systems that are difficult to source in the Global South due to their high commercial cost and cold-chain shipping requirements. Here, we describe the implementation of RNA toehold switch-based sensors using *E. coli* cell lysate-based cell-free protein expression systems, which can be produced locally and reduce the cost of sensors by two orders of magnitude. We then demonstrate that these in-house cell lysates provide sensor performance comparable to commercial PURE cell-free systems. We further optimize use of these lysates with a CRISPRi strategy to enhance the stability of linear DNAs, enabling the direct use of PCR products for fast screening of new designs. As a proof-of-concept, we develop novel toehold sensors for the plant pathogen Potato Virus Y (PVY), which dramatically reduces the yield of this important staple crop. The local implementation of low-cost cell-free toehold sensors could enable biosensing capacity at the regional level and lead to more decentralized models for global surveillance of infectious disease.

## Introduction

Cell-free sensors have emerged as a promising technology for fast and field-deployable detection of nucleic acids (Pardee et al., 2014, 2016; Ma et al., 2018; Takahashi et al., 2018; Verosloff et al., 2019) and chemical pollutants (Didovyk et al., 2017; Jung et al., 2020; Liu et al., 2020; Silverman et al., 2020; Thavarajah et al., 2020). Cell-free toehold RNA sensors (CFTS) have been shown to detect viruses such as Ebola (Pardee et al., 2014), norovirus (Ma et al., 2018), and Zika (ZIKV) (Pardee et al., 2016) at femtomolar concentrations of viral RNA when combined with isothermal RNA amplification (Pardee et al., 2016). Toehold RNA sensors are *de novo* designed genetic structures that can be programmed to detect RNAs of arbitrary sequences (Green et al., 2014). A toehold switch relies on the formation of a hairpin loop at the 5′-end of a reporter gene that structurally prohibits an interaction between the ribosome binding site (RBS) and ribosomes, thereby preventing translation. This structure is released via toehold-mediated strand displacement by an unstructured RNA ‘trigger’ with sequence-specific complementarity to the switch (Green et al., 2014), thus enabling ribosomal-binding and subsequent protein translation. These CFTS can be lyophilized and transported at room temperature for in-field detection of viral RNAs (Pardee et al., 2014, 2016; Ma et al., 2018).

CFTS are implemented in a reconstituted cell-free expression system, known commercially as PURExpress, which consists of a mixture of 34 individually purified components involved in transcription, translation and energy generation (Shimizu et al., 2001). Despite the potential of CFTS for point-of-care deployment, constraints in the accessibility to PURE systems in South America impose significant barriers to locally engineer *de novo* sensors. PURE systems are expensive and proprietary (Laohakunakorn et al., 2020). Furthermore, they need to be imported from the Northern hemisphere under cold-chain shipping, which increases their costs and the risk of activity loss. PURE systems can also be prepared in the laboratory by purifying the components individually or as a single step from a bacterial consortium of strains (Shimizu et al., 2001; Horiya et al., 2017; Villarreal et al., 2018; Lavickova and Maerkl, 2019). Alternatively, cell lysate-based protein expression systems, made from *E. coli* crude extract that do not require purification of particular components, can also be used as a substrate for CFTS (Silverman et al., 2019). Lehr et al. have shown the implementation of CFTS in commercially available cell-free transcription-translation systems (TX-TL) based on cell extracts (Lehr et al., 2019), which are also sourced from the US and shipped under cold chain. Encouragingly, Silverman et al. (2019) and McNerney et al. (2019) have recently shown the successful implementation of CFTS in home-made cell extracts, offering a path for local prototyping *de novo* sensors of interest (McNerney et al., 2019; Silverman et al., 2019).

Cell extracts are cell lysate-based protein expression systems that have been instrumental to molecular biology and biotechnology for decades (Nirenberg and Matthaei, 1961). Cell extracts have also become a versatile tool for engineering complex biomolecular systems outside living cells (Garenne and Noireaux, 2019; Laohakunakorn et al., 2020). Diverse genetic systems have been implemented in cell extracts, e.g. for sensing small molecules (Salehi et al., 2018; Jung et al., 2020; Silverman et al., 2020; Thavarajah et al., 2020), engineering metabolic pathways (Dudley et al., 2016; Grubbe et al., 2020; Swartz, 2020), producing virus-like particles (Bundy et al., 2008; Rustad et al., 2018), and prototyping genetic devices (Siegal-Gaskins et al., 2014; Moore et al., 2017), including RNA circuits (Takahashi et al., 2015), transcriptional cascades (Shin and Noireaux, 2012; Garamella et al., 2016) and oscillators (Niederholtmeyer et al., 2015; Tayar et al., 2017; Yelleswarapu et al., 2018). Various protocols have been crafted to simplify extract preparation and optimize lysate performance, including methods based on cell homogenization (Kim et al., 2006), sonication (Shrestha et al., 2012; Kwon and Jewett, 2015), bead-beating (Sun et al., 2013), enzymatic lysis or flash freezing (Didovyk et al., 2017).

With the goal of making cell-free toehold sensor technology more accessible and affordable, here, we describe the implementation of CFTS in cell extracts supplemented with maltodextrin as an energy source. These CFTS, can detect RNA at similar sensitivity to the PURExpress-based CFTS but for a fraction of the cost. Furthermore, using a CRISPRi strategy for silencing nucleases before harvesting cells, we increased DNA stability, enabling cell-free protein synthesis from linear, PCR-derived templates. These optimized extracts were suitable for fast prototyping of novel toehold sensors directly from linear PCR products using a lacZ gene template. As a proof-of-concept demonstration, we applied our framework to engineer cost-effective CFTS for PVY virus detection, which poses a significant economic threat to local potato farmers and producers of potato seeds.

## Materials and Methods

### Crude extract preparation

BL21 DE3 Star cells (Invitrogen, C601003), BL21dLacGold cells (Didovyk et al., 2017) (Agilent, 230132), or the CRISPR+, CRISPR-derivatives from this work were grown overnight at 37 °C on 2xYT agar plates with the appropriate antibiotics. A single colony was grown in a 5 ml 2xYT culture overnight at 37 °C. The next day, a 50 mL culture was started with a 1:1000 dilution overnight at 37 °C. On the fourth day, a total of 2 liters of culture was started and split in 5 2-liter Erlenmeyer, at an initial OD_600_ 0.05 in 2xYT supplemented with 40 mM phosphate dibasic (Sigma, 94046), 22 mM phosphate monobasic (Sigma, 74092) and 20 g/L D-glucose. This culture was grown at 37 °C with vigorous agitation (230 rpm) until OD_600_ reached 0.6-0.8 (approximately 2.5 hours) and cells were induced with 1 mM IPTG for another 3 hours at 37 °C before harvesting by centrifugation at 5000 g and 4 °C. Pellets were washed twice with cold S30B buffer (5 mM tris acetate pH 8.2, 14 mM magnesium acetate, 60 mM potassium glutamate, and 2 mM dithiothreitol) (Sun et al., 2013). The pellet was then weighted, and a relationship of 0.9 mL of S30B and 5 g of 0.1 mm diameter silica beads (Biospec, 11079101) were added per gram of pellet obtained. 2.5 mL bead-beating tubes and cups were filled with the viscous solution composed of cells and glass beads, without generating bubbles, sealed and beaten for 30 seconds using a homogenizer (MP Biomedicals, FastPrep-24 5G). To remove the beads from extracts, processed samples were placed on the top of a micro-chromatography column (Biorad, 7326204), which was placed into an empty bead-beating tube. The bead-beaten tube attached to the micro-chromatography column and the empty recipient tube was placed into a 15 ml Falcon tube and centrifuged at 6000 g for 5 minutes at 4 °C. Properly beat extracts should appear clear, and two distinct layers should be observed. The supernatant from all tubes was pooled and agitated at 37 °C inside a 5 ml unsealed tube for one hour for runoff. After the run-off reaction, the tubes were placed on ice and re-centrifuged at 6000 g for 10 minutes at 4 °C to pull down the debris generated during the run-off reaction. The supernatant from this centrifugation, i.e. the crude extract, was aliquoted, frozen, and stored at −80 °C until use. A link to the full protocol of crude extract and cell-free reaction preparation can be found here: https://www.protocols.io/view/preparation-of-cell-free-rnapt7-reactions-kz2cx8e.

### Cell-free transcription-translation reaction

The in-house cell-free reactions were composed of 45% crude lysate and 40% reaction buffer. The remaining 15% included DNA input solution chlorophenol red-β-D-galactopyranoside (Sigma, 59767 at final concentration of 0.6 mg/mL), and in the case of the RNA sensing reactions, trigger RNA from in vitro transcription. A typical reaction is composed of 50 mM HEPES pH 8, 1.5 mM ATP and GTP, 1.4 mM CTP and UTP as triphosphate ribonucleotides, 0.2 mg/ml tRNA (Roche, 01109541001), 0.26 mM Coenzyme A, 0.33 mM NAD, 0.756 mM cAMP, 0.0068 mM folinic acid, 0.112 mg/mL spermidine, 2% PEG-8000, 3.4 mM of each of the 20 amino acids (glutamate is also added in excess in potassium and magnesium salts), and 12 mg/ mL maltodextrin and 0.6 mg/mL sodium hexametaphosphate (Sigma, 305553) or 30 mM 3-PGA as an energy source. Cell-free reactions were mounted on ice in a final volume of 10 µL in 1.5 ml Eppendorf tubes and vortexed for 30 seconds before sampling 5µL that were loaded in V-bottom 96 well plates (Corning, CLS3957) or rounded 384 well-plates (Corning, CLS3540). All the reactions were incubated at 29 °C in a Synergy HTX plate reader (BioTek).

### Input DNA preparation

All plasmids used in this work correspond to single transcriptional units (Level 1 plasmids) that were prepared using Golden Gate assembly from Level 0 parts (T7 Promoter, T7 terminator, Toehold Sensors, Triggers, etc.) (Pollak et al., 2019). The level 0 parts were previously prepared by Gibson assembly (Gibson et al., 2009) from PCR linear DNA amplified from gblocks (IDT). All plasmids were commercially sequenced before use. Plasmid DNA input was produced by midi prepping an overnight culture of 200 mL LB with the appropriate strain (Promega, A2492) and cleaned again using PCR cleanup (Promega, A6754). Plasmid DNA inputs were used in a set of concentrations ranging from 0.5 nM to 10 nM. Input linear DNA was produced by PCR and cleaned with PCR cleanup kit (Promega, A6754), quantified and diluted to desired concentrations using ultra-pure water. A list of all the primers and plasmids used in this work is shown in Supplementary Table S1 and Supplementary Table S2, respectively.

### Trigger RNA preparation

PCR products containing the trigger T7 transcriptional unit were cleaned by PCR cleanup kit (Promega, A6754) and 250 ng were processed by *in vitro* transcription using the HiScribe T7 kit (Promega, E2040S) and incubated at 37 °C overnight. The products were then treated with DNAseI for one hour at 37 °C and the enzyme was inactivated by heat at 70°C for 10 minutes. Next, standard RNA clean up steps were performed according to the RNeasy MinElute kit (Qiagen, 74204) before quantification.

### Isothermal NASBA amplification

We used the NASBA lyophilized kit from Life Sciences following the manufacturer’s instructions. Negative controls and RNA dilutions in the range 27 mM to 0.2 fM were prepared in ultra-pure water just before use. The lyophilized reaction buffer containing nucleotide mix (Life Sciences, LRB-10) was reconstituted with DRB-10 diluent (Life Sciences, DRB-10), heated at 41 °C for 5 minutes and kept at room temperature. The initial mixture consisted of 10 µL of the reconstituted reaction buffer, 1 µL RNAsin (Promega, N2111), 380 mM of each DNA primer, 2 µL ultra-pure water and a 1 µL RNA amplicon dilution were assembled at 4 °C and incubated at 65 °C for 2 min, followed by a 10 min annealing at 41 °C. During annealing, the lyophilized enzyme mix, consistent of AMV RT, RNaseH, T7 RNAP, and high molecular weight sugar mix (Life Sciences, LEM-10), was reconstituted with cold D-PDG (Life Sciences, D-PDG-10) and placed on ice. Immediately after annealing, 5 µL of the dissolved enzyme mix was added to the reaction, to a final volume of 20 µL, and the mixture was incubated at 41 °C for 2 h with thermocycler lid at 98 °C. A 10 minute incubation at 70 °C was performed for enzyme inactivation. The amplified product was subsequently cleaned up with the RNeasy MinElute kit (Qiagen, 74204).

### Lyophilization and storage of cell-free reactions

Lyophilization was performed by assembling 9 µL of cell extracts and 8 µL of reaction buffer in a PCR tube. In a separate tube, 3 µL of the corresponding plasmid were mixed with 1 µL of substrate (9 mg/ml), when LacZ reporter was used. Tubes containing the assembled reactions were closed with adhesive aluminum film and punctured with a 16-gauge needle to create a hole. The tubes were placed in a FreeZone 2.5 L Triad Benchtop Freeze Dryer (Labconco), previously pre-chilled at −75 °C (shelf temperature), for overnight freeze-drying of two segments. The first segment was at −75 °C (shelf temperature) with a temperature collector of −80 °C for 12 hours and 0.04 mbar of pressure. The second segment was performed at −20 °C for 4 h with the same pressure. Samples were stored at room temperature in open plastic bags over 30 g of silica inside a closed Tupperware. Rehydration of cell-free reactions was done with 20 µL of a plasmid solution in ultra-pure water. pT7:sfGFP was used at 6 nM or 8 nM final concentration whereas ZIKV Toehold 27 was used at 10 nM final concentration. Trigger 27 RNA was added at a final concentration of 300 nM.

### Design of novel toehold sensors

With the aim of designing novel toehold sensors, we implemented an *in silico* tool called NupackSensors, which is based on an algorithm described by Green et al. (Green et al., 2014) (http://www.github.com/elanibal/NupackSensors). To identify possible triggers from a target sequence, the algorithm first determines all contiguous 36 nucleotide sub-sequences. Each possible trigger defines one specific toehold sensor. After filtering out the sensors that contained stop codons, potential structural defects of each design were calculated, along with a set of thermodynamic parameters (Supplementary Fig. S1, Supplementary Table S3 and S4). The following score function was implemented, and one score value was assigned to each of the possible designs (Ma et al., 2018):

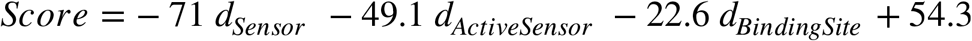

Here, *d*_*Sensor*_ represents the average number of incorrectly paired nucleotides respecting the ideal toehold sensor structure, defined as “…………………….(((((((((((… (((((…………)))))…)))))))))))…………………” (Pardee et al., 2016) in dot-bracket notation (Supplementary Figure S1). Similarly, the parameter *d*_*ActiveSensor*_ represents the average number of incorrectly paired nucleotides respecting the ideal activated sensor structure. *d*_*BindingSite*_ represents the deviations from the perfect single-stranded binding site in the toehold domain, which is essential for the initial trigger binding and subsequent strand-displacement reaction. After assigning one score to each toehold sensor design, they are ranked.

For the PVY toehold sensor design, two RNA target sequences were evaluated in NupackSensors, corresponding to the long and short sequences found to be conserved in several PVY strains (Supplementary Tables S5 and S6). For each of them, four of the highest scores were picked for experimental evaluation. Linear DNA encoding the PVY toeholds were prepared by PCR using specific primers that add the T7 promoter and each toehold sensor sequence on their 5’ tails along with the UNSXR reverse primer. LacZ full length was used as template (Supplementary Table S1 for primer sequences).

### sgRNA design

Sequences of each operon of the genes that are part of the RecBCD complex and the EndI endonuclease were identified in *E*.*coli* K12 strain genome using the web platform RegulonDB (Salgado et al., 2013). sgRNAs were designed according to (Larson et al., 2013)), following these steps: all available 5’->3’PAM sites (CCN) within the nucleotide sequence of the gene of interest were marked, identifying contiguous sites as target candidates; the target candidates were analyzed and selected as the 18-22 bp contiguous to the PAM sequence that are within the first ∼ 156nt of the CDS (i.e. counting from the ATG); sequences that end in C or T were selected so that the sgRNA begins with A or G (corresponding to nucleotide +1 of the sgRNA transcript) since this increases the transcription of the promoter used (Larson et al., 2013; Qi et al., 2013). The designed sgRNA corresponded to the reverse complementary sequences of the chosen target. These steps were performed in ApE open source software (https://jorgensen.biology.utah.edu/wayned/ape/). We designed three sgRNAs that targeted the cistron of *recB* and *recD, recC* and *endA* genes, respectively.

### Stability of PCR-derived linear DNA in cell-free reactions

Linear DNA fragments of 1.5 kb and 500 bp were prepared via PCR for its use as linear DNA input and standards, respectively. Those were purified by wizard gel and PCR clean-up kit (Promega), achieving a final concentration of 200 ng/µL. 600 ng of the 1.5 kb linear DNA were added to 20 µL cell-free reactions and incubated at 29 °C for 0, 30, 60 or 120 minutes and quickly submerged in liquid nitrogen to stop the reaction. Frozen samples were put on ice; and immediately, a volume of binding buffer from PCR clean-up kit (Promega, A9282) was added followed by 2 µL (300 ng) of the 500 bp linear DNA that served as an internal standard for the purification. The samples were then purified following standard supplier’s protocol and eluted with 30 µL of ultra-pure water. Next, 0.3 volumes of loading buffer were added to the purified samples and 10 µL were run on a 1 % agarose gel electrophoresis containing 1X SYBR-Safe for DNA visualization. 1.5 kb and 500 bp DNA fragments were placed in two independent lines along with another line for 1 kb Ladder (NEB, N232). Once bands were clearly identifiable, the gel was photographed on a blue LED light transilluminator with a blue-light filter. The intensity of the 1.5 kb bands in each sample was integrated using ImageJ and normalized with the intensity of the 500 bp band of the same lane.

## RESULTS

### 1. Implementation of an in-house cell-free gene expression system

We first sought to implement of a low-cost and simple protocol for cell-free gene expression. To do this, we combined the use of S12 crude extracts – a simple post lysate processing which does not require ultracentrifugation nor dialysis (Kim et al., 2008) – with a highly concentrated amino acid mixture (Caschera and Noireaux, 2015b) and a cost-effective energy source. This energy source is based on maltodextrin and polyphosphates, which do not require cold-chain transportation (Caschera and Noireaux, 2015a). We characterized our system measuring the dynamics or endpoints of the pT7:sfGFP constitutive expression (Figure 1A). Initially, we prepared four different batches and tested optimum magnesium concentration in the reaction buffer for the cell-free expression of sfGFP. Optimal values ranged between 8.5 nM and 10 nM for all extracts (Supplementary Fig. S2). With the goal of reducing the cost of lysate preparation, we compared maltodextrin and polyphosphates (HMP) to the conventional and more expensive 3-phosphoglyceric acid (3-PGA) energy source. In a side-by-side comparison with 3-PGA, maltodextrin produced a similar initial rate in sfGFP fluorescence (Supplementary Fig. S3). Moreover, the endpoint measurements were significantly higher for the cost-effective maltodextrin-based energy source in each batch tested (Figure 1B). Next, we measured the dynamics of constitutive pT7:sfGFP (Figure 1A) expression in these extracts compared to two RNAP T7-based commercial kits. In comparison to these commercial cell-free systems, our in-house cell lysates generated higher endpoint fluorescent values, although variability was observed between different batches of cell-free extracts (Figure 1C).

**Figure 1:**
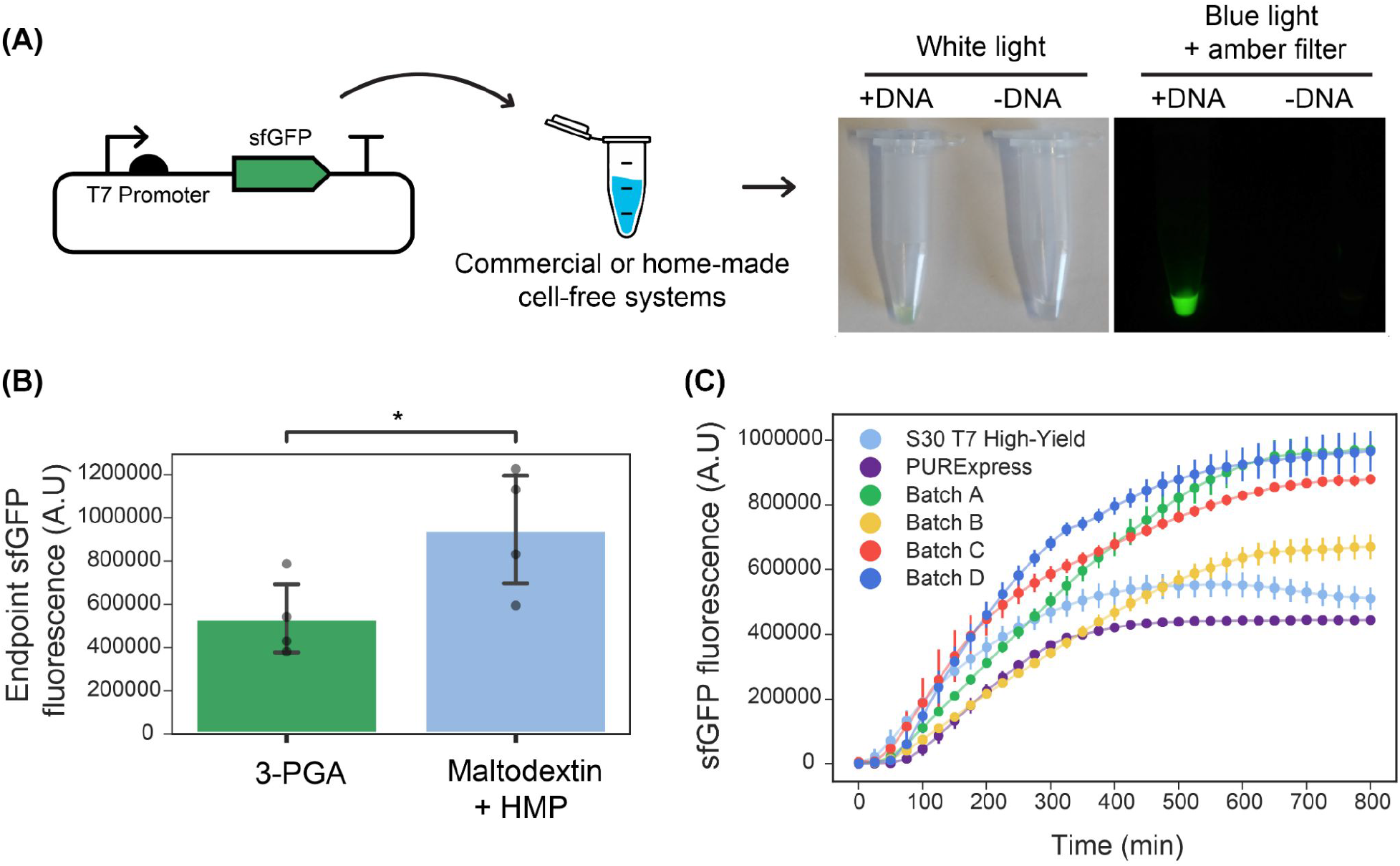
Optimized in-house cell-free reactions compared to commercial alternatives. **(A)** Left: Scheme of the strategy for testing the performance of home-made or commercial cell-free systems using the sfGFP constitutive reporter. Right: Example of the endpoint sfGFP reaction and negative control (without input DNA) in a home-made cell-free system supplemented with maltodextrin energy source. Tubes were photographed under white light or blue light plus an amber filter that allows visualizing the sfGFP fluorescence. (**B)** Endpoint sfGFP fluorescence was measured in four different cell extracts (Batch A, B, C, D) supplemented with either maltodextrin and polyphosphates (light blue) or 3-PGA (green) as energy source. Grey dots represent the arithmetic mean of three measurements performed on each batch, and error bars represent standard deviations of the means of the four batches tested (N=4). T-test for paired measurements was performed and statistically significance was found between the two groups (*p*-value = 0.03, shown by *). Assumptions of the paired t-test were verified using the Shapiro-Wilk test for normality of the differences between energy sources for a given batch (*p*-value = 0.97), and Levene test for homoscedasticity of the 3-PGA and maltodextrin data sets (*p*-value = 0.25). **(C)** sfGFP production dynamics in reactions performed at 29 **°**C using NEB PURExpress and Promega S30 T7 High Yield commercial kits along with four optimized in-house cell-free reactions (Batch A, B, C, D) using maltodextrin and polyphosphates as the energy source. Error bars represent the standard deviations of three independent replicates, dots are centered at the arithmetic mean for each time point.

A cost-breakdown analysis of lab-scale extract production of this optimized cell-free lysate system was performed, comparing reagent prices available to the collaborating institutions in Chile and the UK (Supplementary Tables S7-S13). The cost per reaction (5 µL) in Chile was $0.069 USD for the maltodextrin-based preparation in comparison to $7.80 USD for the locally purchased commercial PURExpress, representing a reduction of two orders of magnitude. In comparison, the home-made preparations were about two times less expensive in the UK than in Chile (Supplementary Fig. S4, Supplementary Table S7). Maltodextrin-based reactions were approximately 20% cheaper than the 3-PGA alternative.

### 2.- Implementation of CFTS in low-cost cell-free reactions

To investigate whether RNA toehold sensors would be functional in the in-house prepared cell extracts, we used Zika virus (ZIKV) toehold sensors already shown to work well in PURExpress (Pardee et al., 2016). We selected ZIKV toehold sensors 8 and 27 and their respective triggers due to their high dynamic range and orthogonality (Pardee et al., 2016). These toehold sensors, controlling full-length LacZ expression, were incubated with the corresponding RNA trigger (400 nM) independently (Fig. 2A). For each sensor, we chose the highest DNA concentration that ensures low-background in PURExpress and used this concentration for comparison with in-house cell-free reactions. Sensor 8 (at 0.8 nM plasmid DNA) behaved similarly in both reaction conditions, achieving a near-maximal response about 120 minutes after induction while maintaining a low signal in the absence of the trigger (Fig. 2B). Sensor 27 (2 nM input DNA), also displayed comparable performance and sequence-specific activation in both systems used (Fig. 2B, 2C). We also compared the performance of the full-length LacZ with LacZ-alpha reporters supplemented with the previously expressed LacZ-omega peptide in commercial and in-house cell-free extracts (Supplementary Fig. S5). Full-length LacZ reporters exhibited higher ON/OFF endpoint absorbance values in both systems tested (Supplementary Fig. S6, far right). This can be explained by the slightly higher OFF state observed on both toehold sensors when using LacZ-alpha as a reporter (Supplementary Fig. S6).

**Figure 2:**
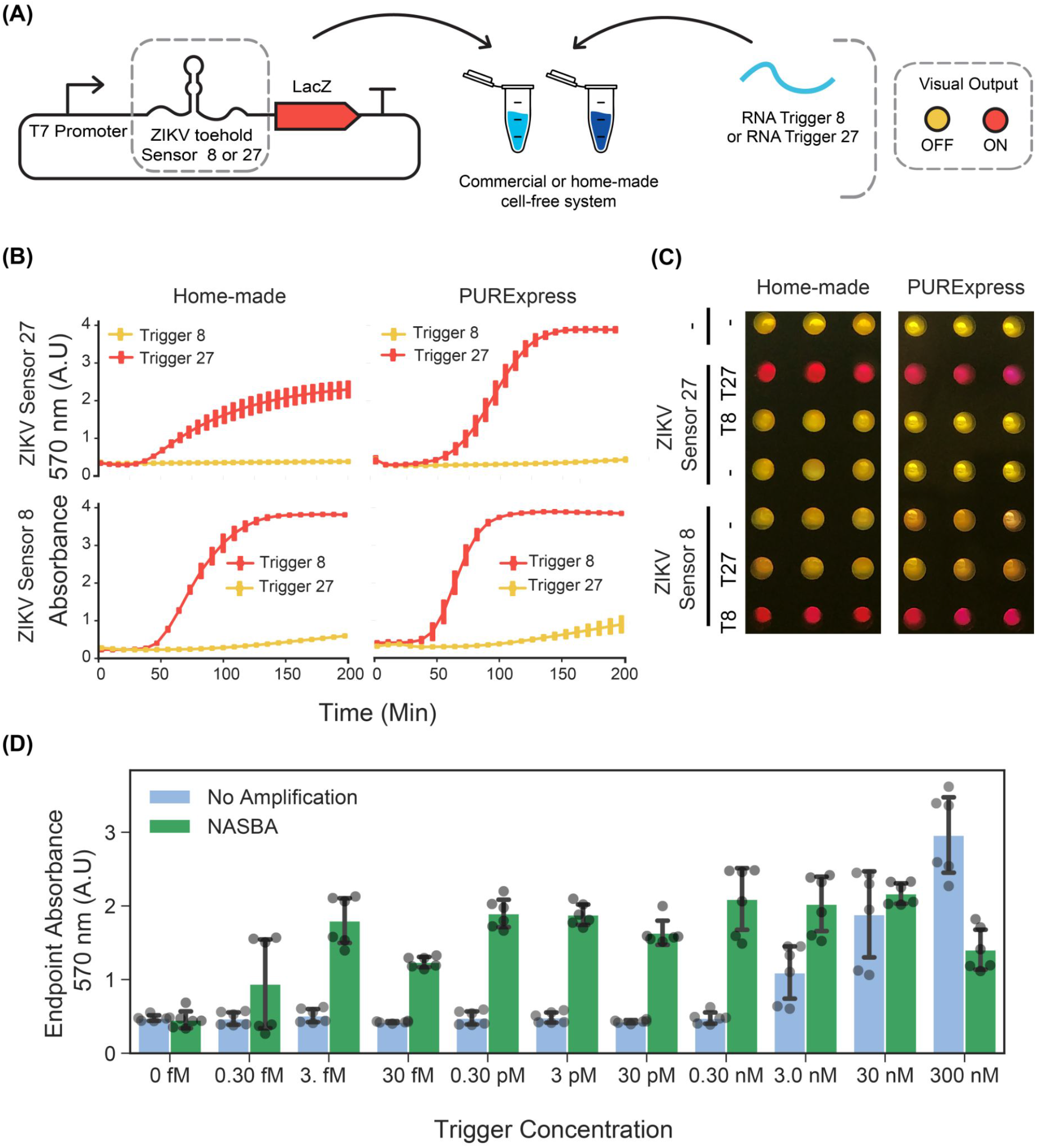
Performance of low-cost ZIKV toehold sensors in low-cost cell-free lysate reactions. **(A)** Schematic representation of toehold-mediated RNA sensing. **(B)** Dynamics of the RNA sensing reactions performed with ZIKV toehold sensor 8 (0.7 nM plasmid DNA) and 27 (2 nM plasmid DNA), regulating the expression of the full-length LacZ in home-made cell extracts and PURExpress cell-free reactions. Error bars represent the standard deviations of three independent experiments, dots are centered at the arithmetic mean for each time point. **(C)** Example of the endpoint visualization of the experiments after four hours of incubation at 29 °C. **(D)** Endpoint measurement of RNA sensing reactions performed with ZIKV sensor 27 and trigger 27 in a range of concentrations with and without NASBA isothermal amplification. Gray dots represent data from six independent measurements performed from two independent NASBA amplifications performed on different days. Black error bars correspond to standard deviations of these six measurements.

Next, we studied the effect of plasmid DNA input concentration on the leaky expression and saturation of sensing reactions. We tested three ZIKV toehold sensors controlling full-length LacZ expression (ZIKV Sensors 8, 27, and 32) (Pardee et al., 2016) in concentrations ranging from 0.5 nM to 10 nM. In all sensors, OFF state signal increased as a function of the input DNA concentration. (Supplementary Fig. S7). These results indicate that input DNA concentration can affect the dynamic range of RNA toehold sensors and that optimal concentrations may be different for each sensor.

To increase the sensitivity of CFTS in home-made cell-free preparations, we performed NASBA amplification of the target RNA before adding it to the reactions containing the sensors (Pardee et al., 2016). While unamplified RNA was detected at concentrations no lower than 3 nM, the NASBA amplification of target RNAs upstream in the workflow permitted the detection of RNA at concentrations as low as 3 fM (Figure 2D), representing a sensitivity increment of six orders of magnitude.

### 3.- Lyophilization and shelf-stability of in-house cell-free reactions

Deployment of RNA sensors in point-of-care settings is facilitated by the lyophilization of cell-free reactions, enabling room temperature transportation and use upon rehydration (Pardee et al., 2016). In order to test whether home-made CFTS can also be freeze-dried and stored at room-temperature, we lyophilized reactions supplemented with pT7:sfGFP plasmid. These reactions were sealed immediately after lyophilization and kept at room temperature for up to 90 days in a closed tupperware containing desiccant and, upon rehydration, compared to a fresh sample for reference. We observed more than 60 % recovery of the endpoint sfGFP expression one week after lyophilization, which decreased to 17 % after 90 days at room temperature (Supplementary Fig. S8). The ZIKV Sensor 27 was also tested in home-made cell extracts after one and seven days of lyophilization. Besides the evident decrease in dynamic range, especially after seven days, these reactions retained the ability to detect its target trigger, while maintaining target specificity (Supplementary Fig. S9). Taken together, these results indicate that the home-made cell-free system is compatible with lyophilization both for constitutive protein expression and for toehold RNA sensing reactions. The efficiency loss during storage at room temperature indicates that the preservation conditions should be addressed for further optimization and the use of cryoprotectants (Karig et al., 2017).

### 4.- Increasing PCR-product stability in home-made cell-free reactions

The use of linear DNA for CFTS offers several advantages compared to plasmid DNA that has to be generated by cloning and then amplified using bacterial cultures prior to use (Yang et al., 1980; Bassett and Rawson, 1983; Thomas Hoffmann et al., 2001; Michel-Reydellet et al., 2005; Seki et al., 2008; Shrestha et al., 2012; Sun et al., 2014; Marshall et al., 2017). Linear DNA can be produced by PCR amplification from a lacZ gene template, enabling the design of toehold structures as primer tails for rapid screening of CFTS libraries. The use of linear dsDNA in cell-free, however, is affected by nuclease-mediated DNA degradation in crude cell extracts. This effect is primarily due to the action of Exonuclease V (Yang et al., 1980; Bassett and Rawson, 1983) (encoded in the RecBCD operon) (Dillingham and Kowalczykowski, 2008) and Endonuclease I (encoded by *end*A), which are the dominant sources of endonuclease activity in *E. coli* (I. R. Lehman et al., 1962). Previous studies have shown an increase in the efficiency of cell-free protein synthesis from PCR products using several approaches, such as supplementing with Exonuclease V inhibitors (Sitaraman et al., 2004; Sun et al., 2014), using modified nucleotides downstream the PCR reaction (Thomas Hoffmann et al., 2001), incorporating competitive DNA strands containing χ-sites (Marshall et al., 2017), adding dsDNA binding proteins (Zhu et al., 2020), depleting Exonuclease V in the crude extracts (Seki et al., 2008), deleting RecBCD with the lambda phage red recombinase (Michel-Reydellet et al., 2005), and generating strains lacking *endA* (Michel-Reydellet et al., 2005; Hong et al., 2015).

To increase the stability of PCR-derived linear DNA, we explored a CRISPRi-based strategy (Larson et al., 2013) to control the genes encoding Exonuclease V and Endonuclease I in cells before harvesting. We designed sgRNAs targeting the 5’ ends of these genes and an IPTG-inducible dCas9 (Supplementary Fig. S10). Once established in the *E. coli* strain of interest, this method is cost-effective as it does not require the supplementation with extra components, or *in vitro* modifications of the PCR product. To test this approach, our crude extracts of each genotype (BL21 DE3 STAR/CRISPRi+, BL21 DE3 STAR/CRISPRi- and BL21 DE3 STAR) were prepared and the stability of a 1.5 Kb linear DNA was quantified over a 2-hour time course using agarose gel electrophoresis (Supplementary Fig. S11). As a control, an off-target sgRNA that does not have a known target in the *E*.*coli* genome (Nuñez et al., 2017) was tested in parallel. The following exponential degradation model was fitted to the data:

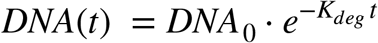

where *DNA*_0_ is the initial concentration of DNA, *DNA*(*t*) is the remaining concentration of DNA at a given time *t*, and *K*_*deg*_ the degradation rate of the DNA. Modelling was undertaken assuming the lower plateau equal to zero, and the ordinary least squares approach was used for parameter estimation. Model screening indicated no statistical evidence of the parameters *DNA*_0_ being different across the data sets evaluated (F-test for extra sum of squares, P=0.9285), finding the best fit value at *DNA*_0_ = 0.9797. Statistical analysis of the *K*_*deg*_ parameters showed significant differences between cell extract genotypes (F-test for extra sum of squares, *p*<0.001). Degradation rate decreased significantly between the CRISPRi-optimized extracts (*K*_*deg*_ = 0.008/*min*) to the two controls (*K*_*deg*_ = 0.022/*min* for the CRISPRi-negative control, and *K*_*deg*_ = 0.040/*min* for the BL21-derived crude extract). Also, the CRISPRi optimized extracts significantly increased protein production capacity from PCR products, leading to a two-fold increase in the sfGFP endpoint signal (Supplementary Fig. S12).

### 5.- Fast Prototyping *de-novo* designed sensors

We prepared CRISPRi-optimized extracts using the BL21-Gold-dLac strain and tested the RNA sensing capability of ZIKV toehold sensors expressed directly from PCR linear products. Using these CRISPRi extracts, both ZIKV sensors, 8 and 27, were able to detect RNA in a sequence-specific manner and behaved similarly to the plasmid derived sensors (Supplementary Fig. S13). Subsequently, we used these CRISPRi-optimized crude extracts to prototype novel RNA sensors for Potato Virus Y (PVY), a virus affecting local agriculture in Chile. First, we selected a conserved sequence from a wide range of PVY strains (Supplementary Fig. S14), which was subsequently generated as a synthetic fragment, transcribed *in vitro* and used for screening potential toehold sensors (Figure 3, Supplementary Table S5 and S6).

**Figure 3:**
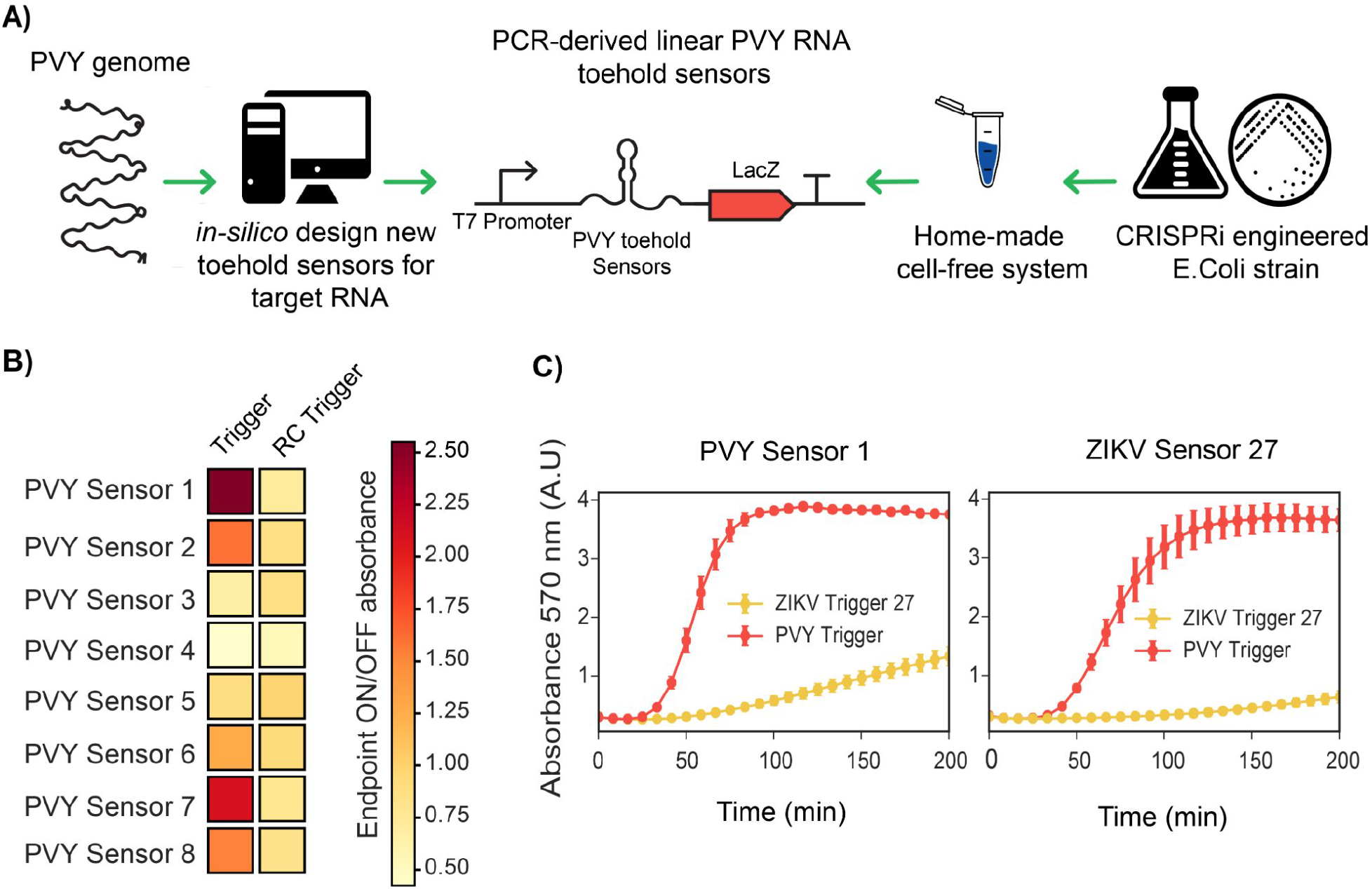
Fast prototyping *de-novo* designed sensors in low-cost optimized cell-free systems. **(A)** Scheme of the global strategy for fast prototyping *de-novo* designed RNA toeholds sensors against synthetic fragments of PVY virus. **(B)** PCR-purified transcriptional units (at 10 nM final concentration) encoding for PVY RNA toehold sensors were incubated with synthetic RNA direct or RNA reverse complementary (RC) (at 300 nM final concentration) and absorbance at 570 nm was measured in plate reader. Endpoint ON/OFF absorbance was calculated at 200 minutes with respect to an untriggered control. Heatmap values correspond to the average of six experimental replicates from two PCR amplifications performed on different days. **(C)** PVY Sensor 1 and ZIKV Sensor 27 encoded in plasmids (at 3.5 nM final concentration) were incubated with ZIKV Trigger 27 RNA or with PVY Trigger RNA (at 480 nM final concentration). Plotted values correspond to arithmetic mean and standard deviation of two independent experiments.

The majority of published toehold sensors have been designed with NUPACK, a software tool for modeling the thermodynamics properties of DNA and RNA molecules (Green et al., 2014; Pardee et al., 2016; Ma et al., 2018). In this work, we have developed an algorithm that leverages NUPACK and can be generally applied to the analysis and design of novel toehold sensors according to ideal designs (Supplementary Fig. S1). Using NupackSensors (see Methods), we analyzed the consensus sequence identified previously and selected eight different toehold RNA sensors for experimental screening. Transcriptional units encoding for each PVY sensor were prepared by PCR amplification and incubated with synthetic RNA direct or reverse complementary (RC) to test their ability to detect RNA in a sequence-specific manner (Figure 3A, Supplementary Figure S15). A control without RNA was used as a baseline in the ON/OFF endpoint calculations. Among the 8 sensors tested, two PVY sensors had ON/OFF endpoint measurements greater than 2 and background level lower than 1.2 (Figure 3B).

In order to validate the fast characterization of PVY toehold sensors using PCR-derived linear DNA, the best scored sensors (PVY Sensor 1 and PVY Sensor 7) were cloned in plasmids and compared side by side with the plasmid version of ZIKV Sensor 27 in a gradient of input DNA concentrations ranging from 1nM to 4 nM (Supplementary Figure S16). Consistent with previous observations (Supplementary Figure S7), input DNA concentrations affect the performance of the toehold sensors, finding best performance of PVY Sensor 7 at a lower concentration than for PVY Sensor 1 whose optimal concentration was between 2 nM to 4 nM. Once a good working concentration was identified, PVY Sensor 1 and ZIKV Sensor 27 were incubated in home-made cell-free reactions (at 3.5 nM final concentration) and supplemented with ZIKV Trigger 27 or PVY RNA (at 480 nM final concentration). PVY Sensor 1 displayed sequence-specific activation comparable with ZIKV Sensor 27 (Figure 3C). These results showed that the fast-prototyping pipeline presented here, followed by an optimization of input DNA concentration, is a suitable strategy for identifying novel RNA toehold sensors of good performance.

## Discussion

The increasing frequency of infectious disease outbreaks and the long-standing impact of viral pathogens on agriculture and livestock highlight the need for more efficient, cost-effective and ubiquitous surveillance of viruses. Cell-free toehold sensors have emerged as a promising technology to address this challenge (Pardee et al., 2014, 2016; Ma et al., 2018). However, in Latin America the high cost and shipping constraints of the commercial *in vitro* transcription and translation systems has limited the local production and distributed development of new sensors. These constraints could lead to centralized production models and fragile technological dependencies prone to failure, as evidenced recently by the disrupted supply chain of diagnostic reagents for SARS-CoV-2 testing (The COVID-19 testing debacle, 2020; John Nkengasong). These shortages highlight the relevance of biotechnological tools that can be shared, produced and deployed locally at a lower cost and without the restrictions of intellectual property (Kellner et al., 2020; Mascuch et al., 2020; Thomas G.W. et al., 2020; JOGL (Just One Gigant Lab)-Open COVID19 Initiative, 2020; Reclone, 2020). In-house prepared cell-free toehold sensors, implemented in cell lysates or home-made PURE systems (Shimizu et al., 2001; Villarreal et al., 2018; Lavickova and Maerkl, 2019), could contribute to the decentralization of this technology and its adaptation to local needs.

Here, we demonstrate the implementation and rapid prototyping of novel sensors using low-cost, locally produced cell-free lysates. As our cost analysis showed, we estimate that this approach could enable the manufacture of bespoke diagnostic tests to meet local needs (for instance in Chile the cost was $0.069 USD/test, considering a 5 µL reaction). While the RNA amplification step still requires commercial inputs, such reactions can also be made in-house (J. Compton, 1991; Lauren M. Aufdembrink et al., 2020). Future work will focus on the in-house preparation of NASBA from locally purified, and IP-free RNAseH, T7 RNAP, pyrophosphatase, and reverse transcriptase enzymes (Lauren M. Aufdembrink et al., 2020; Reclone, 2020; Matute et al., 2021).

Future lines of research will also address the cost efficiency of crude extract preparation. 45-60% of the maltodextrin-based reaction cost relates to growing, inducing, and processing cells, where IPTG (used for induction) and the micro-chromatography columns (used for separation of the cell extract from the glass beads) add up to more than one third of the cost per reaction (Supplementary Figure S4). Therefore, sonication (Shrestha et al., 2012), or other column-free methods for disrupting the cells should be evaluated, along with the use of alternatives to the relatively expensive IPTG such as use of the autolysis strain (Didovyk et al., 2017) or the autoinduction media formulation (Levine et al., 2019).

A key aspect of cell-free biology is the possibility of freeze-drying genetic systems into pellets or paper-based reactions that can be transported at room temperature and deployed in the field. Here, we have tested the performance of toehold sensors in lyophilized cell extracts stored at room temperature for up to seven days. The decay of the activity after lyophilization highlights the need for further optimization of the storage conditions. Modifications of the storage atmosphere conditions either by replacing air with an inert gas, such as nitrogen or argon, in addition to the use of vacuum sealing, oxygen absorbers, and desiccants (Jung et al., 2020), remains to be explored.

## Author Contributions

FF conceived the project, supervised the research, assisted in preparing the figures and writing the manuscript.

AA designed and performed the experiments, analyzed the results, prepared the figures and wrote the first draft of the manuscript.

ND assisted the coding writing, data analysis and statistics.

JP, FGC, CG, TM conceived and performed experiments, analyzed the results.

KP, JM, JH, ND supervised the research, assisted in writing the manuscript.

## Funding

AA was founded by CONICYT scholarship N° 21140714.

FF, AA, TM were funded by ANID - Millennium Science Initiative Program - ICN17_022, Fondo de Desarrollo de Áreas Prioritarias - Center for Genome Regulation (ANID/FONDAP/ 15090007) and ICGEB CRP/CHL19-01

FF, AA, JP Proyecto Investigación Interdisciplinaria 2018 VRI

## Acknowledgements

We thank Dr. Jonathan Dhalin (Technical University of Denmark) and Dr. Vincent Noireaux (University of Minnesota) for providing positive controls and examples for cell-free protein synthesis reactions at early stages of the project.

We thank the Pardee lab (University of Toronto) for welcoming Anibal during his internship. We thank the Federici lab (Pontificia Universidad Católica de Chile) for comments and feedback on the manuscript.

