## Supplementary information for "Decentralizing cell-free RNA sensing with the use of low-cost cell extracts"

### *Supplementary Material*

#### 1 Supplementary Figures and Tables

##### 1.1 Supplementary Figures

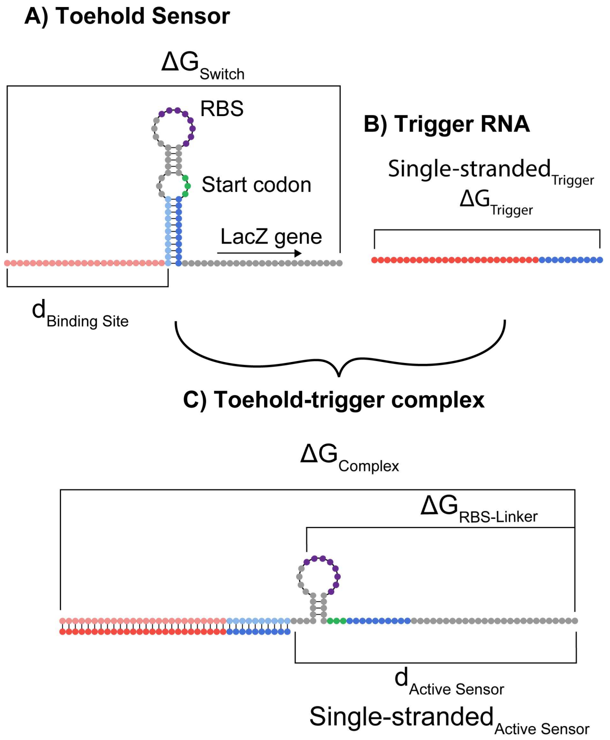


**Supplementary Figure S1: Ideal structures of (A) toehold sensor RNA, (B) trigger RNA, and the (C) toehold-trigger complex.** Trigger RNA (B) is complementary to the binding site of the toehold sensor (A) and its first 10 nucleotides of the hairpin structure. Strand-displacement reactions between trigger RNA and toehold sensor causes the formation of the toehold-trigger complex (C). Each circle represents one ribonucleotide. Complementary sequences are shown in the same color but different intensities. The sequence used to define each parameter is also shown.

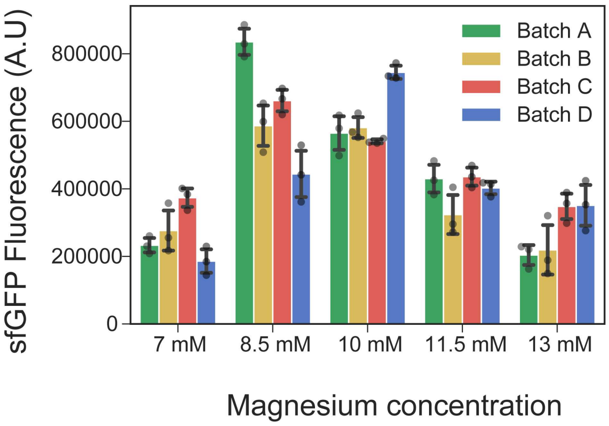


**Supplementary Figure S2: Magnesium concentration optimization for the in-house cell-free preparations**. Endpoint sfGFP fluorescence measurements in cell-free reactions performed in four different crude extracts (Batch A, B, C, D) supplemented with maltodextrin energy buffers containing a range of magnesium concentrations. Optimal levels for magnesium were found between 8.5-10 mM for all extracts. Data points of three independent experiments are shown in light grey circles while black error bars represent the standard deviations.

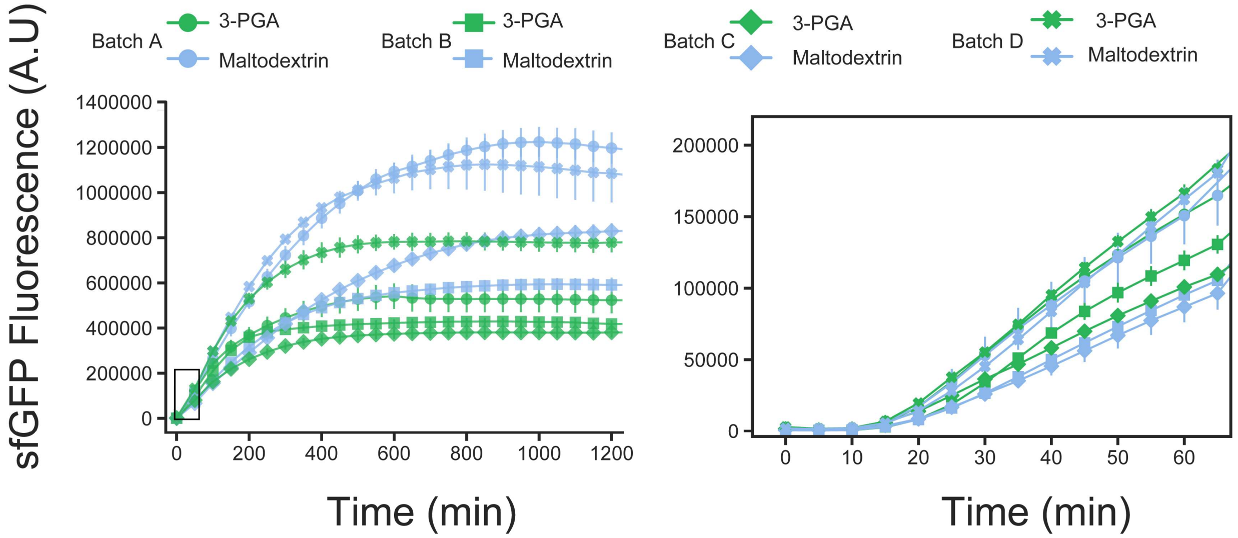


**Supplementary Figure S3: Dynamics of sfGFP production from constitutive expression in cell-free reactions supplemented with 3-PGA or Maltodextrin-based energy sources.** Constitutive sfGFP production on batches A, B, C and D supplemented with maltodextrin (light blue) or 3-PGA (green) energy buffers. Right, magnified inlet rectangle showing sfGFP production during the first hour of reaction. Data points are centered at the average of three measurements per batch, error bars represent the standard deviations.

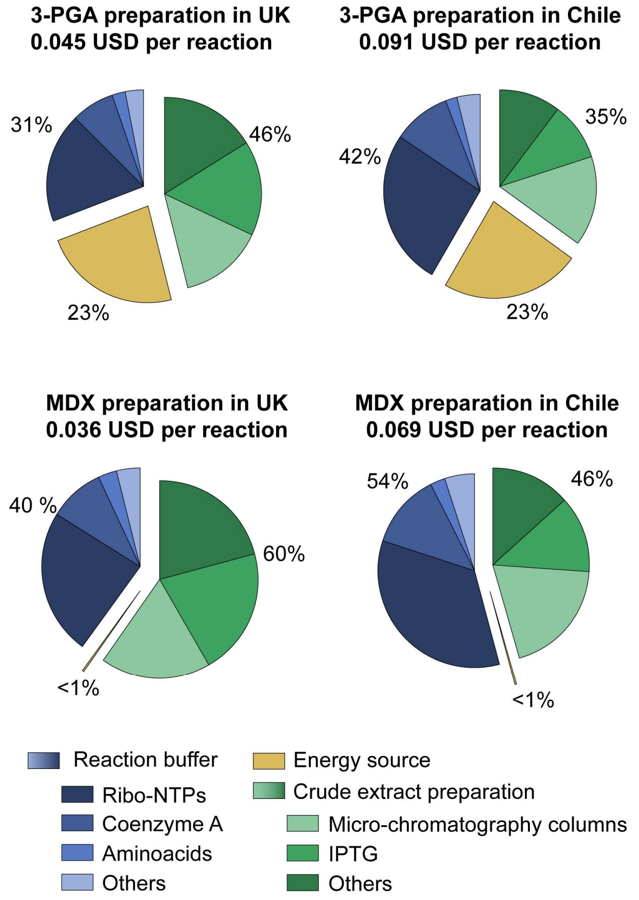


**Supplementary Figure S4: Cost comparison of reactions prepared in Chile and in the UK using 3-PGA or maltodextrin (MDX) preparation protocols.** The fraction of the costs representing energy source is shown in yellow, crude extract preparation reagents in green and reaction buffer in blue. We assumed 1 liter of culture yields 6 g of biomass, needing 8 micro-chromatography columns to be processed, representing 3.75 ml of crude extract per liter of culture.

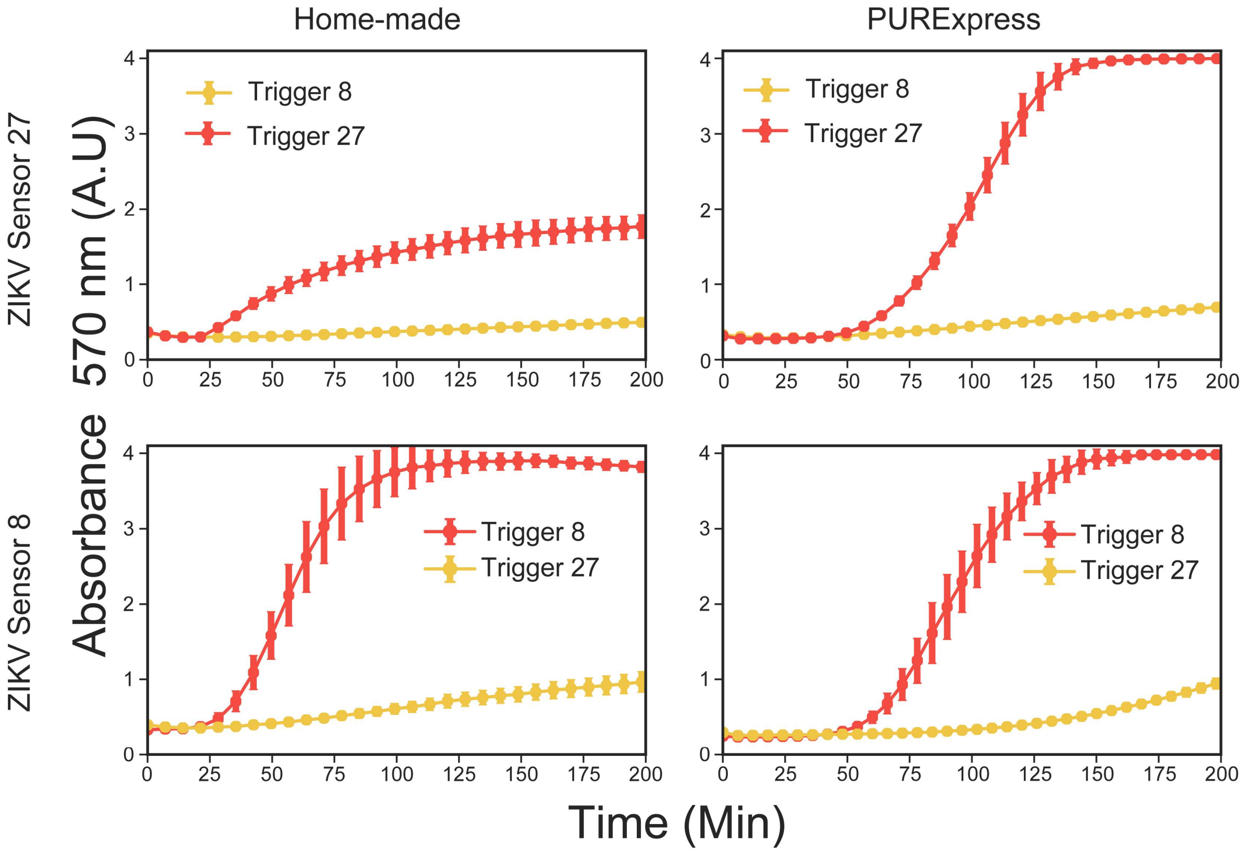


**Supplementary Figure S5: Comparison of RNA sensing reactions performed on in-house cell-free preparations and commercial PURExpress using LacZ-Alpha as reporter.** Dynamics of RNA sensing reactions performed with the ZIKV toehold sensor 8 (0.7nM plasmid DNA) and 27 (2 nM plasmid DNA), regulating the expression of the LacZ-Alpha. Reactions were performed in home-made cell-free preparations (left) or commercial PURExpress (right). These reactions were supplemented with the pre-synthesized LacZ-omega peptide to a final concentration of about 2 µM. Error bars represent the standard deviations of three independent replicates, dots are centered at the arithmetic mean for each time point.

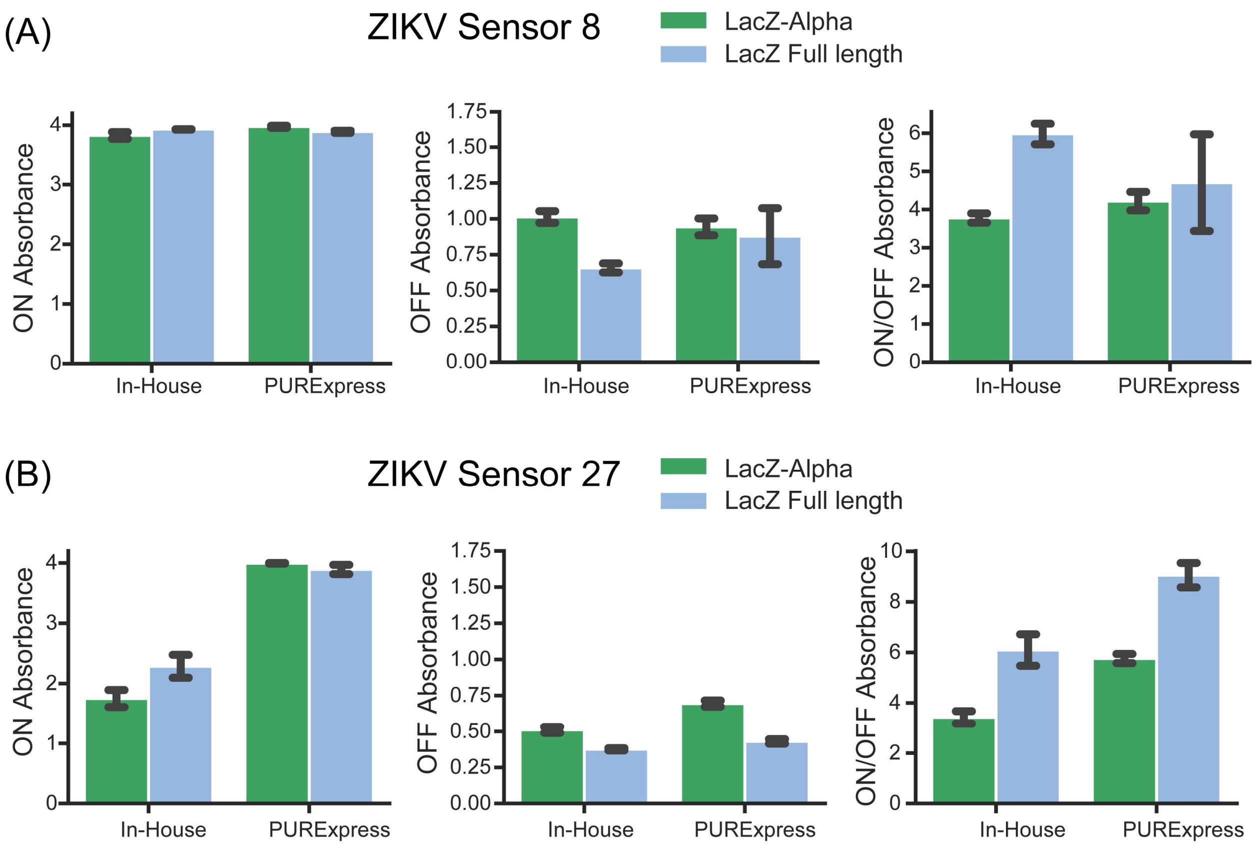


**Supplementary Figure S6: Performance comparison of the ZIKV toehold sensor 8 and 27 using LacZ-Alpha as reporter.** Reporters LacZ-Alpha (green) and full-length LacZ (light blue) were compared at endpoint absorbance (at 570 nm) for the trigger-activated (ON), untriggered (OFF), and ON/OFF measurements of ZIKV Sensor 8 (a) or ZIKV Sensor 27 (b) after 200 minutes of incubation at 29 °C. Sensor 8 was used at 0.7 nM plasmid DNA concentration, Sensor 27 at 2 nM and trigger RNAs at 270 nM. Error bars represent the standard deviations of three independent replicates, dots are centered at the arithmetic mean for each time point.

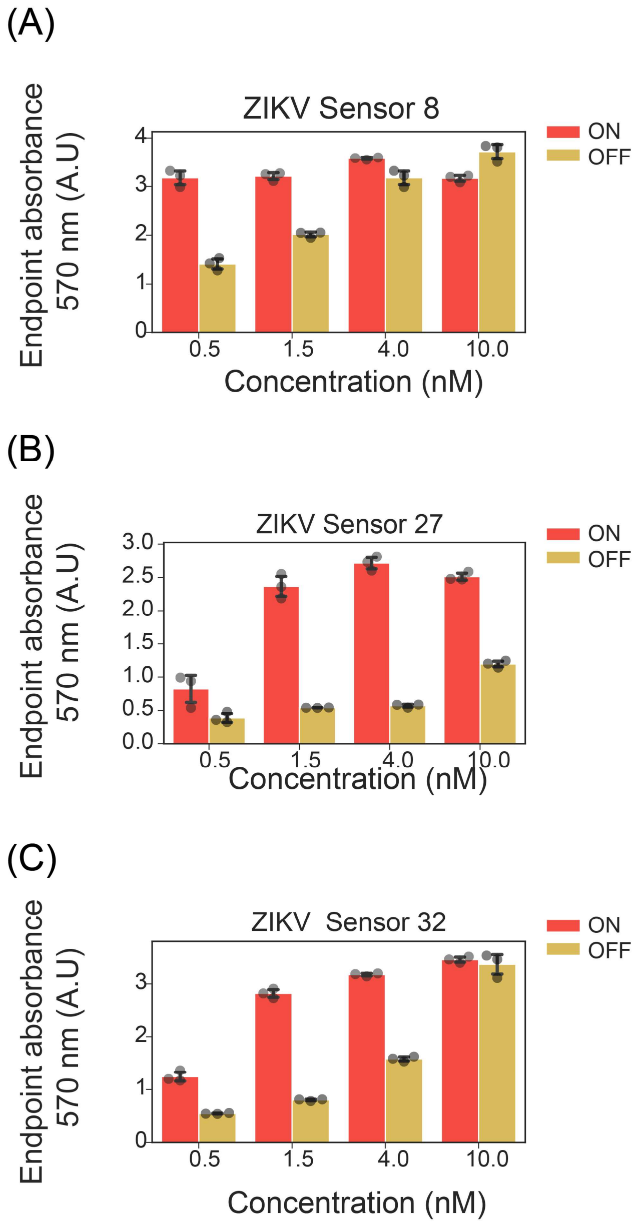


**Supplementary Figure S7: Plasmid concentration effect on the response dynamic range of ZIKV toehold sensors 8, 28, and 32.** Endpoint absorbance of RNA sensing reactions were performed with the ZIKV toehold Sensor 8 **(A)**, Sensor 27 **(B)**, and Sensor 32 **(C)** at different concentrations of starting plasmid DNA, ranging from 0.5 nM to 10 nM. Reactions in the presence of the corresponding trigger are shown in red (ON), while reactions without a trigger are shown in yellow (OFF). Grey dots represent three experimental replicates for each concentration while standard deviation of these is shown in black error bars.

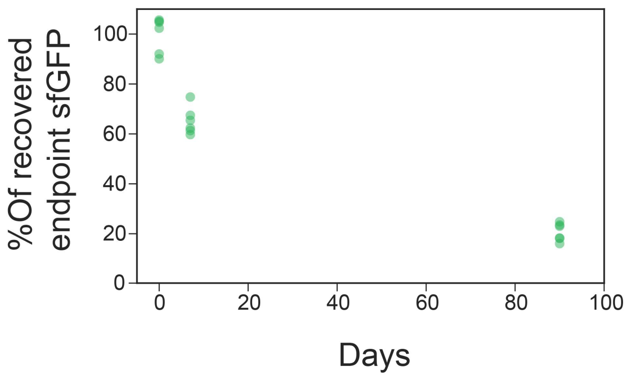


**Supplementary Figure S8: Shelf stability of the lyophilized cell-free reactions over three months.** Reactions were stored for up to ninety days at room temperature after lyophilization. More than 60% recovery was observed after seven days, and 17% recovery after ninety days at room temperature.

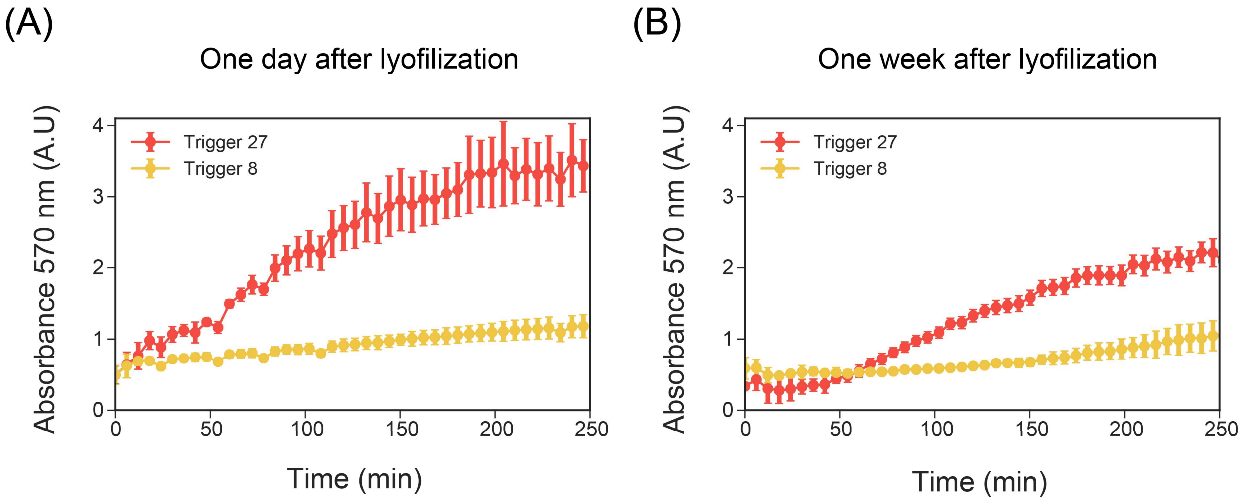


**Supplementary Figure S9**: **Supplementary Figure S8: Performance of ZIKV Sensor 27 after lyophilization**. ZIKV toehold sensor 27 was evaluated one **(A)** and seven days **(B)** after lyophilization and storage at room temperature.

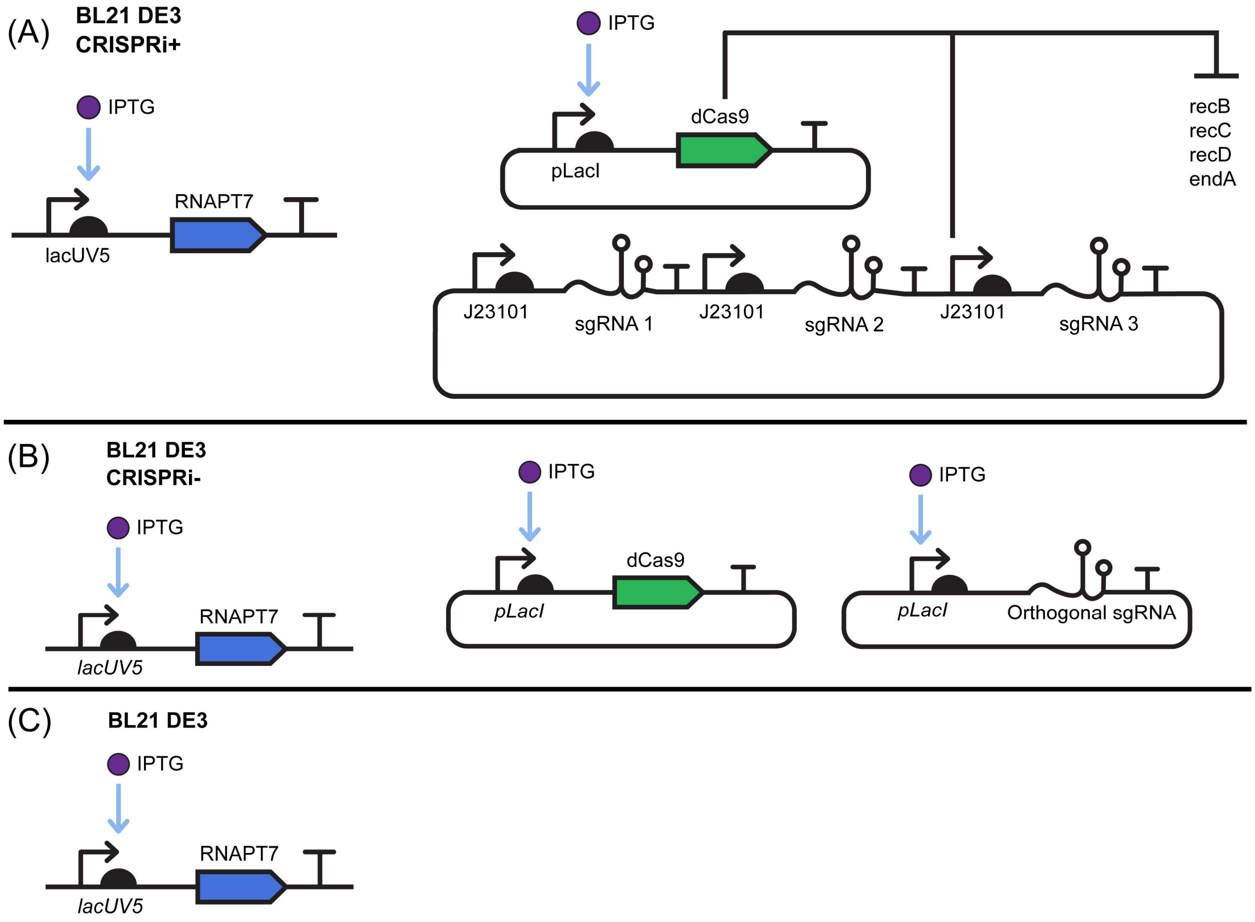


**Supplementary Figure S10: Schematic representation of CRISPRi genetic circuits used for nuclease silencing. (A)** In the CRISPRi+ strain, IPTG induces the expression of RNAPT7 and dCas9, and a complementary plasmid enables the constitutive expression of three sgRNAs that target recB, recC, recD, and endA genes. **(B)** In the control CRISPRi- strain, IPTG induces the expression of dCas9 in addition to the RNAPT7, a complementary plasmid supports the expression of the "non-targeting" sgRNA that does not have any target in the genome. **(C)** In the basal genotype strain, BL21 DE3 STAR, IPTG only induces the production of RNAPT7 before harvesting the cells.

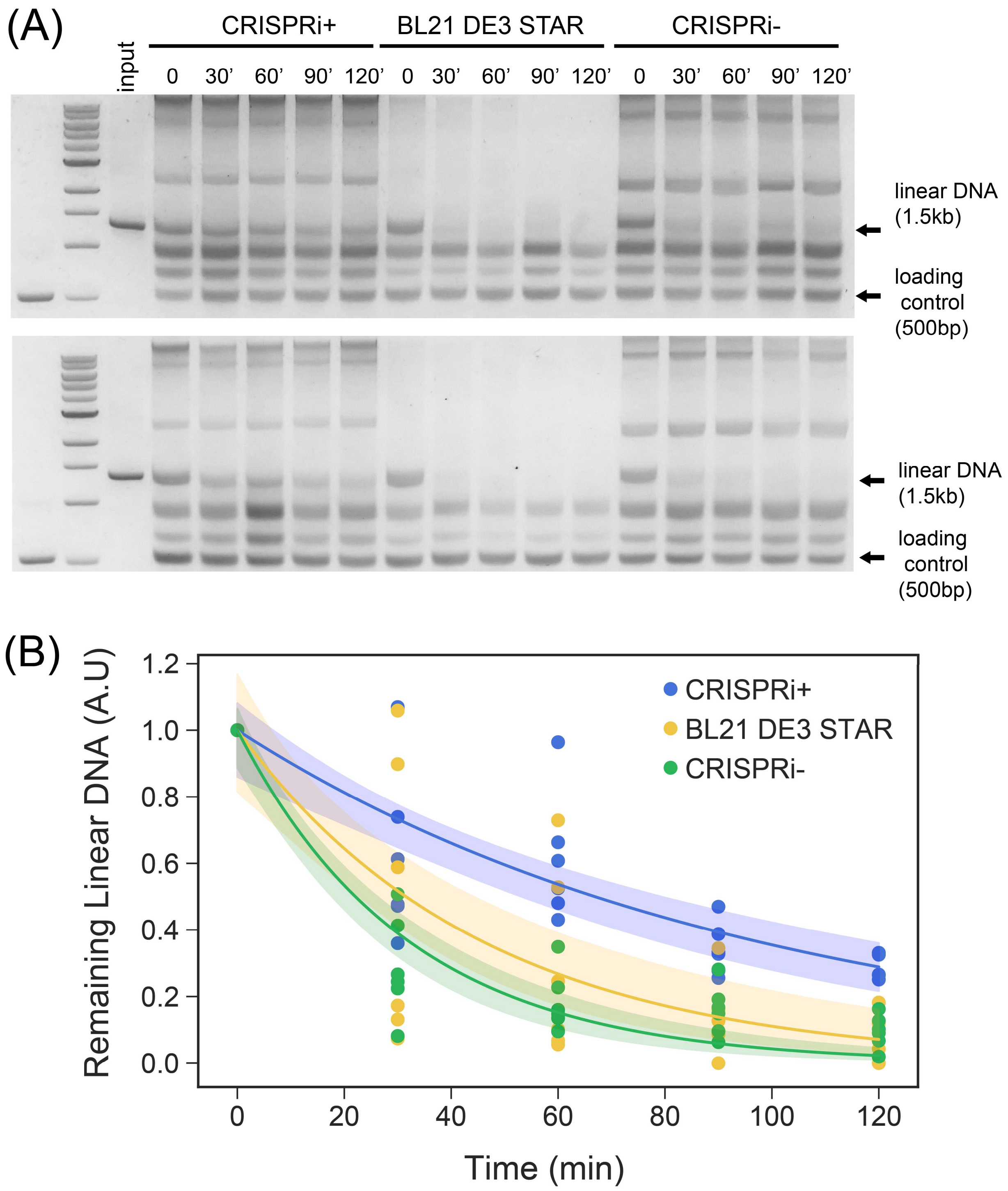


**Supplementary Figure S11: Increased stability of PCR-derived linear DNA in CRISPRi optimized cell-free extracts. (A)** Example of two independent experiments measuring the stability of a linear product of 1.5 kb that was incubated in cell-free reactions during a range of 120 minutes. Agarose gels were used for resolving the linear DNA product of interest and integration of its signal, 500 pb was used as a loading control for quantification. Experiments were performed in three different genotypes: CRISPR+, CRISPR- and BL21 DE3 STAR. (**B)**

Exponential DNA degradation models for each genotype evaluated. Each dot represents a single measurement for a given batch of each genotype. A total of six batches per genotype were tested. The colored areas indicate 95% confidence intervals for the model, which was approximated by sampling values of $DNA_0$ and $K_{deg}$ from a multivariate Gaussian distribution with mean and covariance identified from inferences performed per genotype. F-test for extra sum of squares was performed comparing two models: a simple model where all the data is explained by a common $K_{deg}$ parameter, or the unrestricted model where the data is explained with an independent $K_{deg}$ parameter per genotype, finding statistical significance to reject the simple model ($p<0.001$). According to the unrestricted model, the degradation rate $K_{deg}$, decreased significantly in the CRISPRi-optimized extracts ($K_{deg}$ = 0.010/min; $R^2$=0.73) with respect to the controls ($K_{deg}$= 0.021/min; $R^2$=0.70 for the CRISPRi-negative control, and $K_{deg}$= 0.031/min; $R^2$=0.90 for the BL21-derived crude extract).

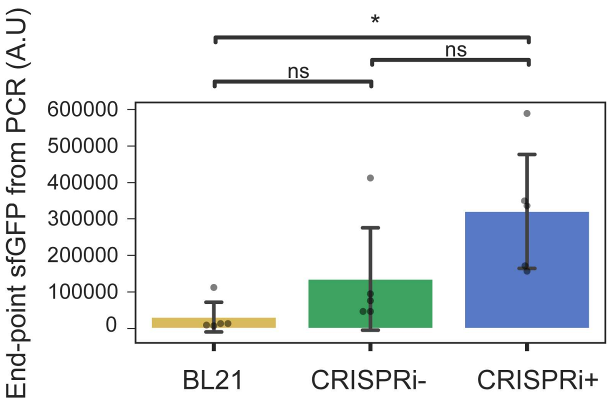


**Supplementary Figure S12: Increased endpoint sfGFP production from constitutive expression of linear DNA in CRISPRi optimized cell-free extracts.** Endpoint sfGFP expression from PCR products was measured in five cell-free extracts from each of the three different genotypes: CRISPRi-optimized (BL21 DE3 STAR/CRISPRi+), negative control for CRISPRi (BL21 DE3 STAR/CRISPRi-), and the basal genotype (BL21 DE3 STAR). Gray dots represent the arithmetic mean of three to six sfGFP measurements performed on a single batch. Black error bars represent standard deviation of the mean values computed for the five different batches of each genotype. Variance homogeneity of the different data sets was tested with Levene test ($p$-value 0.35) while normality of the data was tested with Shapiro-Wilk test ($p$-value 0.0084). Therefore, non-parametric one-way Kruskal-Wallis test was performed obtaining $p$-value of 0.012. Pairwise Mann-Whitney-Wilcoxon test for independent multiple comparisons with Bonferroni corrections were performed as a post hoc test identifying significant differences between the CRISPRi+ and the BL21 ($p$-value=0.036).

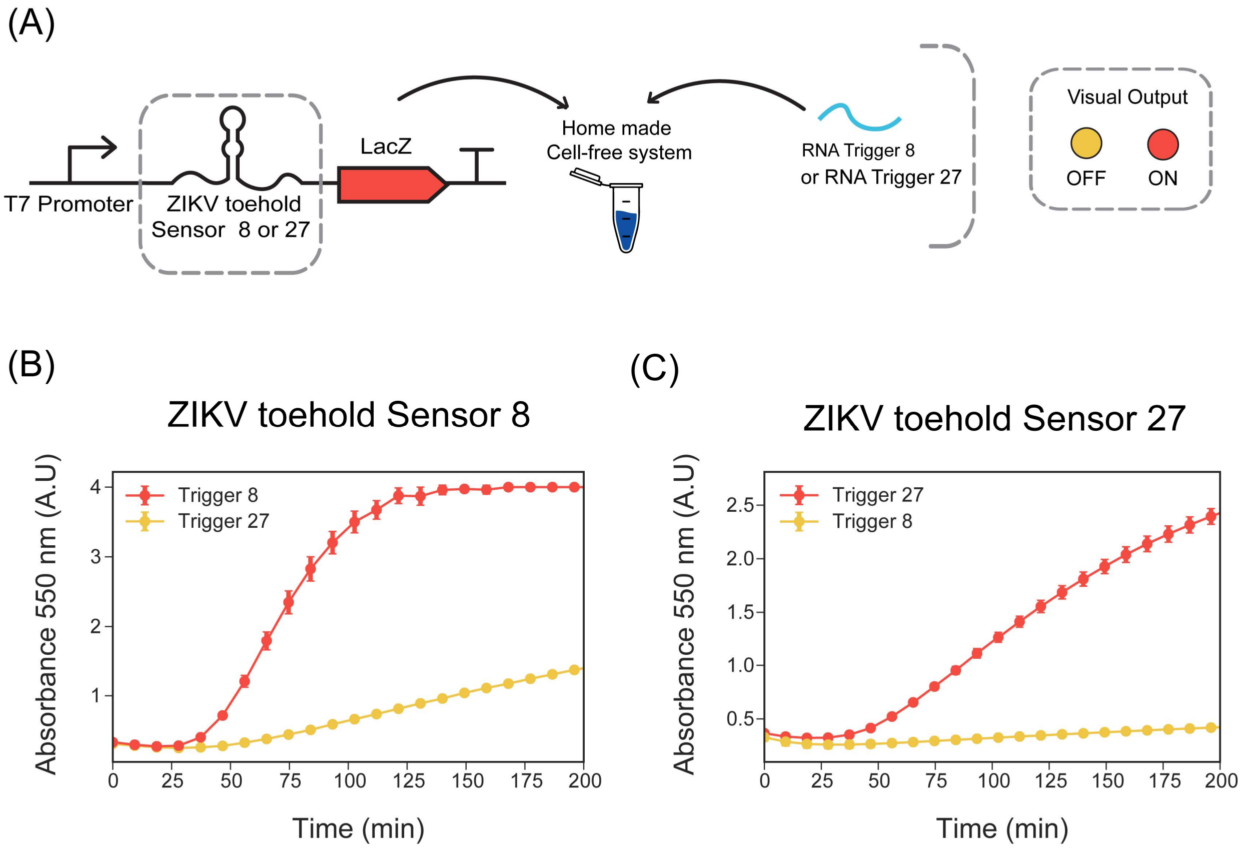


**Supplementary Figure S13**: **Performance of RNA sensing reactions using PCR-derived linear DNA encoding for ZIKV toehold sensors**. (A) Scheme of the experiment performed. Dynamical profile of RNA sensing reactions using ZIKV toehold Sensor 8 **(B)** and Sensor 27 **(C)**. Linear DNA encoding both sensors were tested at 10 nM in the reaction and were incubated with 300 nM of the trigger RNA. Dots are centered at the arithmetic mean of three experimental replicates while error bars represent standard deviations.

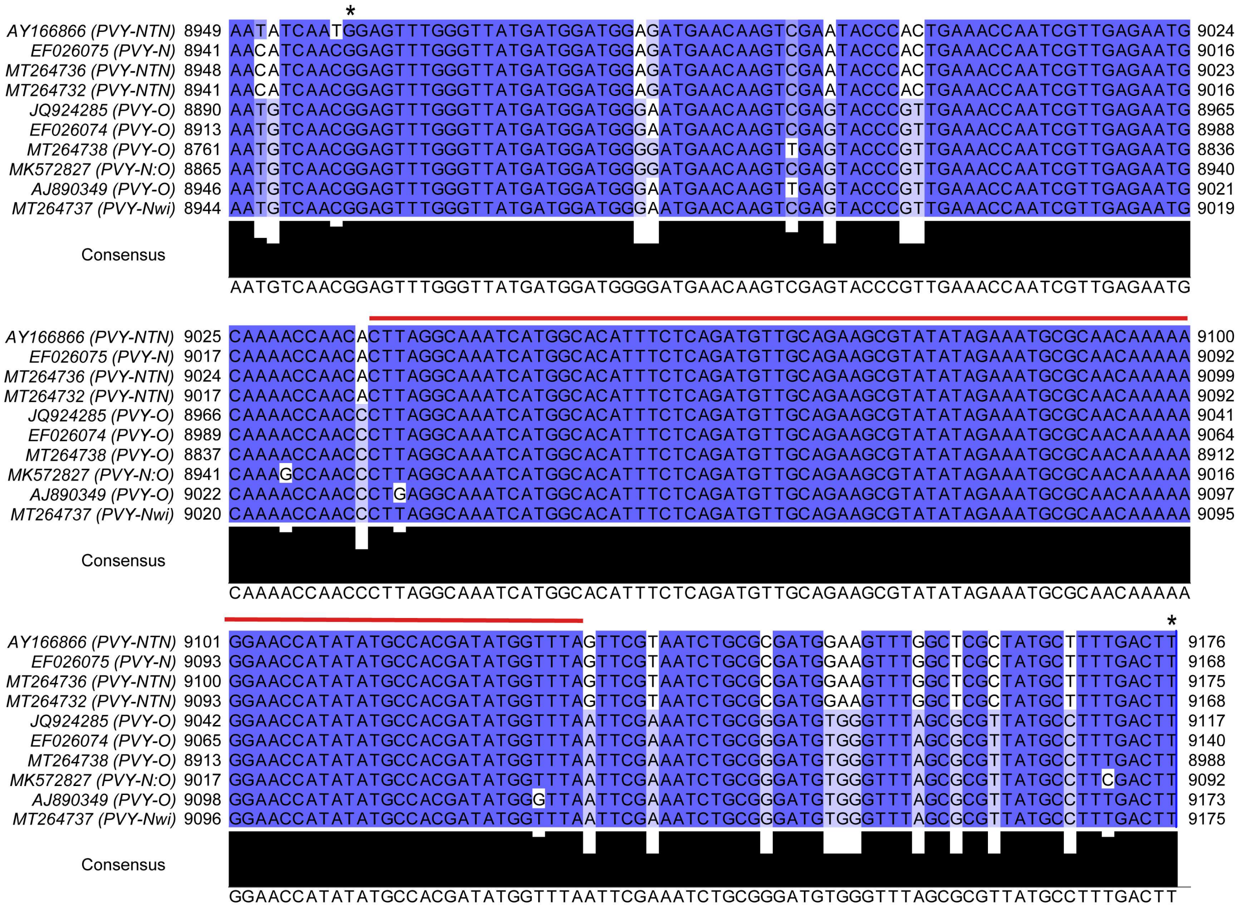

**Supplementary Figure S14: Conserved region across ten PVY genomes.** GenBank entries corresponding to PVY viruses of different viral strains: EF026075.1 (PVY-NTN), AY166866.1 (PVY-NTN), MT264736.1 (PVY-NTN), MT264732.1, (PVY-N), MT264737.1 (PVY-NWi), MK572827.1 (PVY-N:O), MT264738.1 (PVY-O), JQ924285.1 (PVY-O), EF026074.1 (PVY-O), AJ890349.1 (PVY-O). The consensus sequence of the NTN genotypes contained between (*) marks was named "Long" and was used for screening PVY Toehold sensors S1 to S4 and, while the consensus region highlighted in red line was named "Short" and was used in the screening the toehold S5 to S8, it contains the longest string of 100% conservation across all the genomes analyzed. Numbers at each sides of the alignment represent the nucleotide position of the first and last nucleotide in that alignment relative to each GenBank entry. Blue color gradient represents nucleotide consensus identity relative to each position of the alignment where the solid blue shows 90-100% consensus identity, and light blue 60% consensus.

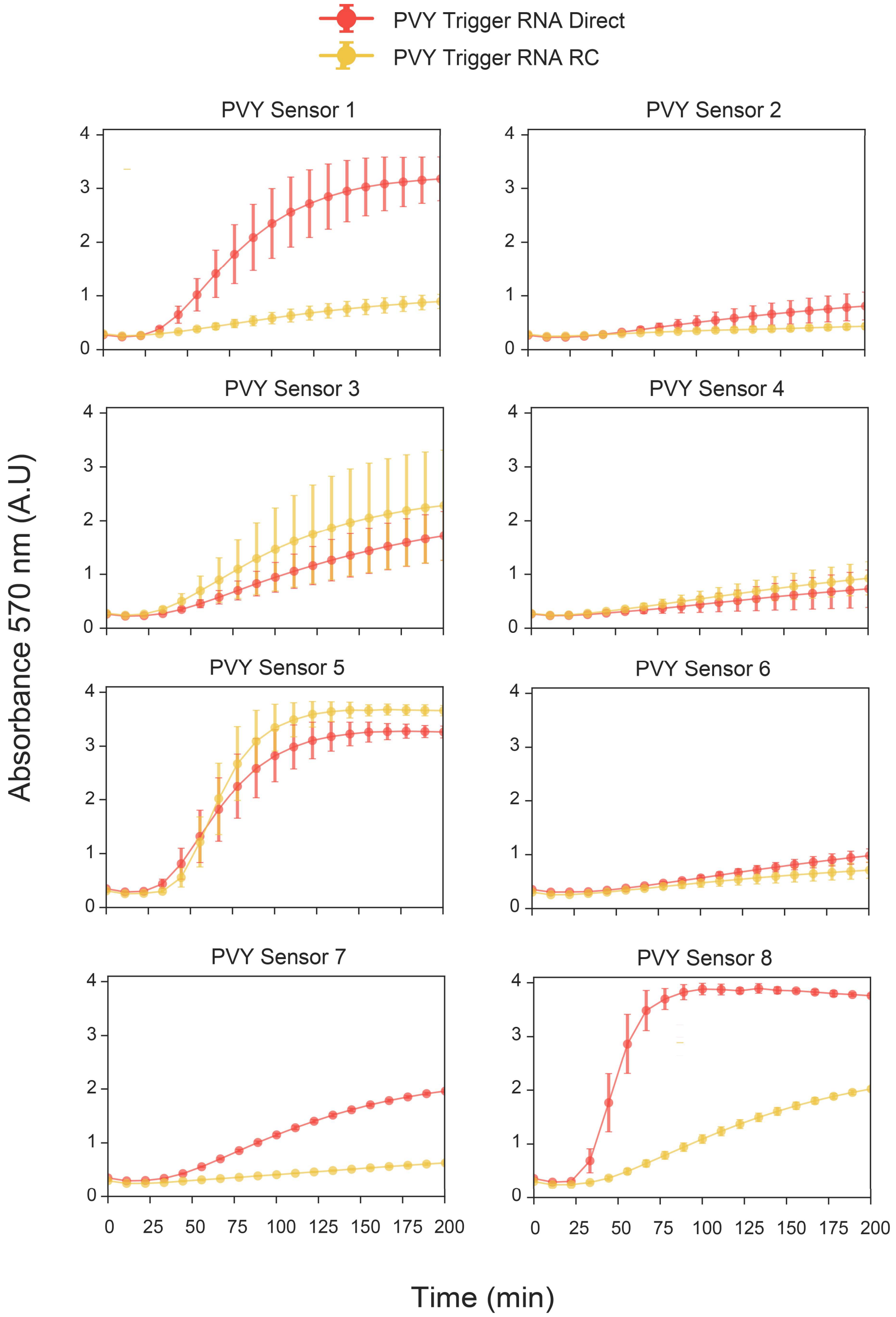


**Supplementary Figure S15: RNA sensing reactions of the linear PVY Toehold sensors 1 to 8 performed in the CRISPRi optimized cell-free extracts.** PVY toehold sensors 1 to 4 were triggered using PVY consensus sequence "Long" in direct (red) or as a control its reverse complementary sequence (yellow). PVY toehold sensors 5 to 8 were triggered using "Short" consensus sequence in direct (red) or as a control, its reverse complementary sequence (yellow). For each plot, dots are centered at the mean of six experimental replicates from two independent PCR amplification. Error bars represent standard deviations of these six measurements.

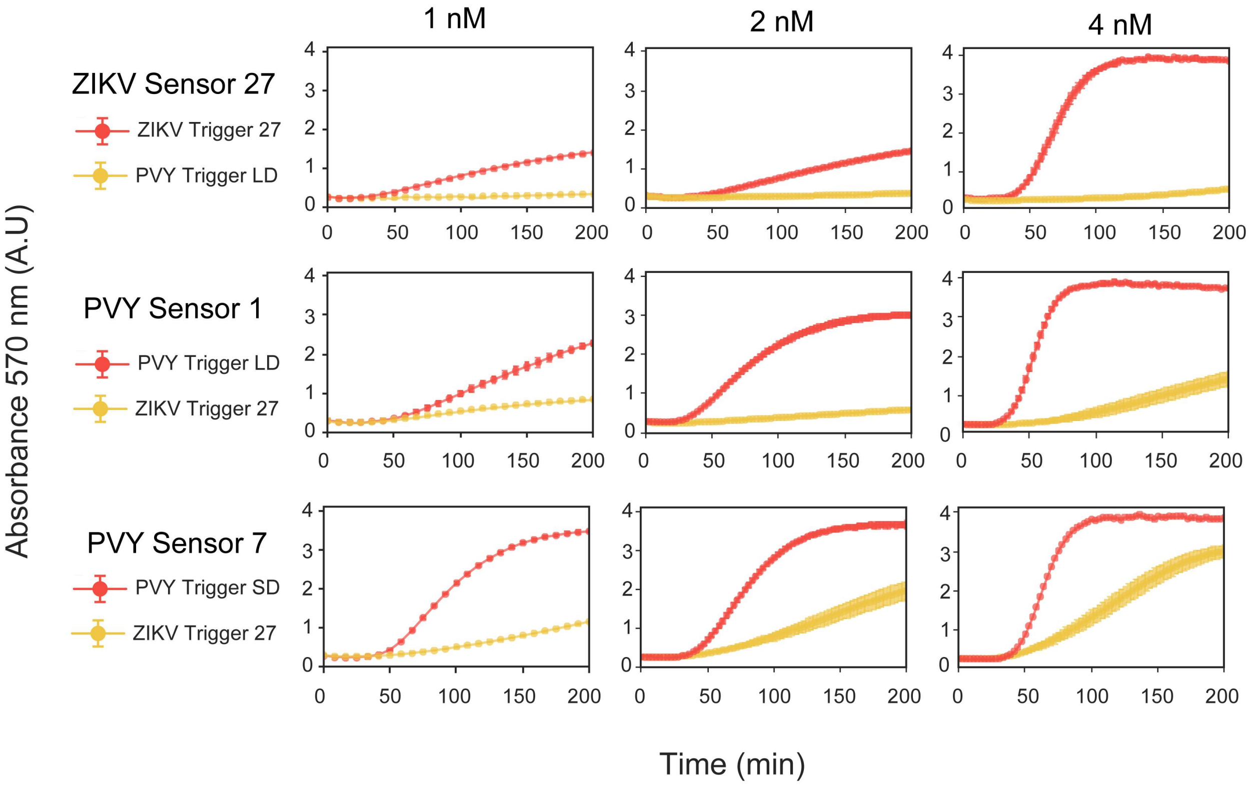


**Supplementary Figure S16: RNA sensing reactions of PVY Toehold Sensors 1 and 7 and ZIKV Sensor 27 at different concentrations of input plasmid DNA.** ZIKV 27, and PVY sensors 1 and 7 encoded in plasmids were evaluated at 1 nM, 2 nM and 4 nM input DNA. PVY Sensor 1 was triggered with PVY Trigger Short Direct RNA (SD, in red) while PVY Sensor 7 was triggered with PVY Trigger Long Direct RNA (LD, in red) dots are centered in the mean of three experimental replicates while error bars represent standard deviations.

#### 1.2 Supplementary Tables

| Primer Name | Sequence | References and notes |
|---|---|---|
| PVY-S1-Fw | CGTTTCTACGGTAGCCGGGCGCTAATACGACTCACTATAGGGCGCAGATTACGAACTAAACCATATCGTGGCATATATGGACTTTAGAACAGAGGAGATAAAGATGATATATGCCACACCTGGCGGCAGCGCAAGAAGAATGACCATGATTACGGATTCACTGGCC | This work. Used for generation of linear PVY toehold sensor S1 |
| PVY-S2-Fw | CGTTTCTACGGTAGCCGGGCGCTAATACGACTCACTATAGGGTCATAAAAGTCAAAAGCATAGCGAGCCAAACTTCCAGGACTTTAGAACAGAGGAGATAAAGATGTGGAAGTTTGGACCTGGCGGCAGCGCAAGAAGAATGACCATGATTACGGATTCACTGGCC | This work. Used for generation of linear PVY toehold sensor S2 |
| PVY-S3-Fw | CGTTTCTACGGTAGCCGGGCGCTAATACGACTCACTATAGGGCGCGCAGATTACGAACTAAACCATATCGTGGCATATGGACTTTAGAACAGAGGAGATAAAGATGATATGCCACGAACCTGGCGGCAGCGCAAGAAGAATGACCATGATTACGGATTCACTGGCC | This work. Used for generation of linear PVY toehold sensor S3 |
| PVY-S4-Fw | CGTTTCTACGGTAGCCGGGCGCTAATACGACTCACTATAGGGTCCATCGCGCAGATTACGAACTAAACCATATCGTGGGGACTTTAGAACAGAGGAGATAAAGATGCCACGATATGGACCTGGCGGCAGCGCAAGAAGAATGACCATGATTACGGATTCACTGGCC | This work. Used for generation of linear PVY toehold sensor S4 |
| PVY-S5-Fw | CGTTTCTACGGTAGCCGGGCGCTAATACGACTCACTATAGGGTGGCATATATGGTTCCTTTTTGTTGCGCATTTCTATGGACTTTAGAACAGAGGAGATAAAGATGATAGAAATGCGACCTGGCGGCAGCGCAAGAAGAATGACCATGATTACGGATTCACTGGCC | This work. Used for generation of linear PVY toehold sensor S5 |
| PVY-S6-Fw | CGTTTCTACGGTAGCCGGGCGCTAATACGACTCACTATAGGGTCGTGGCATATATGGTTCCTTTTTGTTGCGCATTTCGGACTTTAGAACAGAGGAGATAAAGATGGAAATGCGCAAACCTGGCGGCAGCGCAAGAAGAATGACCATGATTACGGATTCACTGGCC | This work. Used for generation of linear PVY toehold sensor S6 |
| PVY-S7-Fw | CGTTTCTACGGTAGCCGGGCGCTAATACGACTCACTATAGGGATACGCTTCTGCAACATCTGAGAAATGTGCCATGATGGACTTTAGAACAGAGGAGATAAAGATGATCATGGCACAACCTGGCGGCAGCGCAAGAAGAATGACCATGATTACGGATTCACTGGCCG | This work. Used for generation of linear PVY toehold sensor S7 |

| | | |
|---|---|---|
| PVY-S8-Fw | CGTTTCTACGGTAGCCGGGCGCTAATACGACTCACTATAGGGTAAACCATATCGTGGCATATATGGTTCCTTTTTGTTGGACTTTAGAACAGAGGAGATAAAGATGAACAAAAAGGAACCTGGCGGCAGCGCAAGAAGAATGACCATGATTACGGATTCACTGGCC | This work. Used for generation of linear PVY toehold sensor S8 |
| U1F | CATTACTCGCATCCATTCTCAGGCTGTCTCGTCTCGTCTC | Used for generation of linear sfGFP and ZIKV Sensor 27 and Sensor 8. Adapted from Torella *et al.*, (2014) |
| U2F | GCTGGGAGTTCGTAGACGGAAACAAACGCAGAATCCAAGC | Used for and PCR amplification of Trigger 8 and 27. Adapted from Torella *et al.*, (2014) |
| UXR | GGTGGAAGGGCTCGGAGTTGTGGTAATCTATGTATCCTGG | Used for generation of linear sfGFP, ZIKV toehold sensors, and triggers 8 and 27. Adapted from Torella *et al.*, (2014) |

**Supplementary Table S1: List of primers used in this work.**

| Plasmid name | Resistance | Notes and references |
|---|---|---|
| pT7_sfGFP | Tetracycline | Allows RNAPT7- mediated expression of sfGFP. |
| pT7_LacZ | Tetracycline | Allows RNAPT7- mediated expression of LacZ. |
| pT7_T8 | Tetracycline | Allows RNAPT7- mediated expression of Trigger 8. Adapted from Pardee *et al.,* (2016). |
| pT7_T27 | Tetracycline | Allows RNAPT7- mediated expression of Trigger 27. Adapted from Pardee *et al.,* (2016). |
| pDM_T7_HisLacZomega | Ampicillin | Allows RNAPT7- mediated expression of the LacZ-Omega peptide. Original from Ma *et al.,*(2018). |
| 12X_S8A_LacZ | Tetracycline | Allows RNAPT7- mediated expression of the ZIKV toehold sensor 8 controlling the expression of LacZ full-length. Adapted from Pardee *et al.,* (2016). |
| 12X_S8A_LacZα | Tetracycline | Allows RNAPT7- mediated expression of the ZIKV toehold sensor 8 controlling the expression of LacZα. Adapted from Pardee *et al.,* (2016). |
| 12X_S27B_LacZ | Tetracycline | Allows RNAPT7- mediated expression of the ZIKV toehold sensor 27 controlling the expression of LacZ full-length. Adapted from Pardee *et al.,* (2016). |
| 12X_S27B_LacZα | Tetracycline | Allows RNAPT7- mediated expression of the ZIKV toehold sensor 27 controlling the expression of LacZα. Adapted from Pardee *et al.,* (2016). |

| | | |
|---|---|---|
| 12X_S32B_LacZ | Tetracycline | Allows RNAPT7- mediated expression of the ZIKV toehold sensor 32 controlling the expression of LacZ full-length. Adapted from Pardee *et al.,* (2016). |
| pT7_ dpCas9 | Ampicillin | Allows RNAPT7- mediated expression of dpCas9. Original from from Nuñez et al.,(2017). |
| ZZ | Kanamycin | Allows the expression of a non-targeting sgRNA, driven by *plac* promoter. Original from from Nuñez et al.,(2017). |
| DD | Kanamycin | Allows the constitutive expression of three sgRNAs driven by three individual J23119 promoters targeting *endA,recB,recC* and *recD.* |
| 12X_PVY_Sensor 1 | Tetracycline | Allows RNAPT7- mediated expression of the PVY toehold sensor 1 controlling the expression of LacZ full-length |
| 12X_PVY_Sensor 7 | Tetracycline | Allows RNAPT7- mediated expression of the PVY toehold sensor 1 controlling the expression of LacZ full-length |

**Supplementary Table S2: List of plasmids used in this work.**

| Parameter | Definition | Notes |
| --- | --- | --- |
| $\Delta G_{Switch}$ | The minimum free energy secondary structure of the toehold sensor. | Calculated at 29°C using the *mfe* function in NUPACK. Values are represented in Kcal/mol units. |
| $\Delta G_{Trigger}$ | The minimum free energy secondary structure of the trigger RNA. | Calculated at 29°C using the *mfe* function in NUPACK. Values are represented in Kcal/mol units. |
| $\Delta G_{Complex}$ | The minimum free energy secondary structure of the activated complex of the toehold sensor with the trigger RNA. | Calculated at 29°C using the *mfe* function in NUPACK. Values are represented in Kcal/mol units. |
| Net $\Delta G_{Complex\ formation}$ | Net ΔG of the toehold sensor and trigger RNA interaction. | Net $\Delta G_{Complex\ formation} = \Delta G_{Complex} - (\Delta G_{Trigger} + \Delta G_{Switch})$ |
| Single-stranded$_{Trigger}$ | The average probability of the nucleotides in the trigger RNA to be unpaired | Calculated at 29°C using the *pairs* function in NUPACK. |
| $\Delta G_{RBS\text{-}Linker}$ | The minimum free energy secondary structure of the sequence between the first nucleotide of the RBS until the end of the toehold sensor. | Calculated at 29°C using the *mfe* function in NUPACK, considering the toehold sensor positions 49 until 96. Values are represented in Kcal/mol units. |
| $d_{Sensor}$ | The average number of incorrectly paired nucleotides in the equilibrium respecting the ideal sensor structure. | Calculated at 29°C using the *complexdefect* function in NUPACK. Ideal structure for Sensor:"........................(((((((((((...(((((............)))))...))))))))))).....................""

| | | |
|---|---|---|
| $d_{Complex}$ | The average number of incorrectly paired nucleotides in the equilibrium respecting the ideal activated toehold-trigger complex. | Calculated at 29°C using the *complexdefect* function in NUPACK. Ideal activated Stoehold-trigger complex:"((((((((((((((((((((((((((((((...(((((............))))).........................+))))))))))))))))))))))))))))))" |
| Single-stranded$_{Active\ Sensor}$ | The average probability of the nucleotides in the active toehold sensor RNA to be unpaired | Calculated at 29°C using the *pairs* function in NUPACK, considering toehod sensor positions 37 until 96. |
| $d_{Binding\ Site}$ | The average number of incorrectly paired nucleotides in the equilibrium respecting the ideal activated trigger binding site. | Calculated at 29°C using the *complexdefect* function in NUPACK, considering toehold sensor sequence from 1 until position 26. |
| $d_{Active\ Sensor}$ | The average number of incorrectly paired nucleotides in the equilibrium respecting the ideal activated toehold sensor | Calculated at 29°C using the *complexdefect* function in NUPACK, considering the sequence of the toehold sensor positions 37 until 96. Ideal structure of Active Sensor:"...(((((............)))))..................................." |
| Score | Metric for scoring toehold sensors design on the *in silico* screening | |

**Supplementary Table S3: Definition of each parameter evaluated over the PVY toehold sensors.**

| Parameter | Values on each toehold design | | | | | | | |
|---|---|---|---|---|---|---|---|---|
| | **PVY S1** | **PVY S2** | **PVY S3** | **PVY S4** | **PVY S5** | **PVY S6** | **PVY S7** | **PVY S8** |
| $\Delta G_{Switch}$ | -25.956 | -28.435 | -29.439 | -30.848 | -26.858 | -29.131 | -31.706 | -32.721 |
| $\Delta G_{Trigger}$ | -7.521 | -8.885 | -7.983 | -6.89 | -2.515 | -3.026 | -7.164 | -9.389 |
| $\Delta G_{Complex}$ | -76.29 | -76.421 | -80.014 | -82.659 | -74.895 | -82.597 | -80.723 | -70.906 |
| Net $\Delta G_{Complex\ formation}$ | -61.248 | -58.651 | -64.048 | -68.879 | -69.865 | -76.545 | -66.395 | -52.128 |
| Single $Streadness_{Trigger}$ | 0.52 | 0.421 | 0.507 | 0.515 | 0.789 | 0.769 | 0.533 | 0.599 |
| $\Delta G_{RBS\text{-}Linker}$ | -8.406 | -8.491 | -8.469 | -8.19 | -6.92 | -10.684 | -8.918 | -5.833 |
| $d_{Sensor}$ | 0.1632 | 0.1667 | 0.2122 | 0.1894 | 0.2229 | 0.2124 | 0.2267 | 0.2328 |
| $d_{Complex}$ | 0.1585 | 0.2089 | 0.1571 | 0.1573 | 0.1554 | 0.1557 | 0.2177 | 0.1322 |
| Single $Streadness_{Active\ Sensor}$ | 0.3408 | 0.3614 | 0.3473 | 0.3601 | 0.34 | 0.3712 | 0.3541 | 0.3009 |
| $d_{Binding\ Site}$ | 0.1335 | 0.1617 | 0.135 | 0.2539 | 0.296 | 0.3029 | 0.3688 | 0.5387 |
| $d_{Active\ Sensor}$ | 0.3209 | 0.4519 | 0.318 | 0.3398 | 0.3153 | 0.3354 | 0.4521 | 0.2695 |
| Score | 24.71 | 17.23 | 22.14 | 19.1 | 17.3 | 16.32 | 8.34 | 12.88 |

**Supplementary Table S4: Values of thermodynamic parameters for each PVY toehold sensor.**

| Description | Sequence |
|---|---|
| DNA used for RNAPT7 *In vitro* transcription of RNA Direct Short | **GATCGATCTCGATCCCGCGAAATTAATACGACTCACTATAGGG**CTTAGGCAAATCATGGCA CATTTCTCAGATGTTGCAGAAGCGTATATAGAAATGCGCAACAAAAAGGAACCATATATGCCA CGATATGGTTTA |
| Direct RNA Short | CUUAGGCAAAUCAUGGCACAUUUCUCAGAUGUUGCAGAAGCGUAUAUAGAAAUGCGCAACA AAAAGGAACCAUAUAUGC CACGAUAUGGUUUA |
| DNA used for RNAPT7 *In vitro* transcription of RNA RC Short | **GATCGATCTCGATCCCGCGAAATTAATACGACTCACTATAGGG** TAAACCATATCGTGGCATATATGGTTCCTTTTTGTTGCGCATTTCTATATACGCTTCTGCAACAT CTGAGAAATGTGCCATGATTTGCCTAAG |
| RNA RC Short | UAAACCAUAUCGUGGCAUAUAUGGUUCCUUUUUGUUGCGCAUUUCUAUAUACGCUUCUGCA ACAUCUGAGAAAUGUGCCAUGAUUUGCCUAAG |

**Supplementary Table S5: DNA and RNA sequences used for screening PVY sensors 1 to 4.** The bold sequence represents the extended T7 promoter used for the generation of each RNA via *in vitro* transcription. RNA sequences were used as input in the NupackSensors algorithm.

| Description | Sequence |
|---|---|
| DNA used for RNAPT7 *In vitro* transcription of RNA Direct Long | **GATCGATCTCGATCCCGCGAAATTAATACGACTCACTATAGGG**GAGTTTGGGTTATGATGG<br>ATGGAGATGAACAAGTCGAATACCCACTGAAACCAATCGTTGAGAATGCAAAACCAACACTT<br>AGGCAAATCATGGCACATTTCTCAGATGTTGCAGAAGCGTATATAGAAATGCGCAACAAAAAG<br>GAACCATATATGCCACGATATGGTTTAGTTCGTAATCTGCGCGATGGAAGTTTGGCTCGCTATG<br>CTTTTGACTTTTATGAAGTT |
| Direct RNA Long | GAGUUUGGGUUAUGAUGGAUGGAGAUGAACAAGUCGAAUACCCACUGAAACCAAUCGUUG<br>AGAAUGCAAAACCAACACUUAGGCAAAUCAUGGCACAUUUCUCAGAUGUUGCAGAAGCGUA<br>UAUAGAAAUGCGCAACAAAAAGGAACCAUAUAUGCCACGAUAUGGUUUAGUUCGUAAUCU<br>GCGCGAUGGAAGUUUGGCUCGCUAUGCUUUUGACUUUUAUGAAGUU |
| DNA used for RNAPT7 *In vitro* transcription of RNA RC Long | **GATCGATCTCGATCCCGCGAAATTAATACGACTCACTATAGGG**AACTTCATAAAAGTCAAA<br>AGCATAGCGAGCCAAACTTCCATCGCGCAGATTACGAACTAAACCATATCGTGGCATATATGG<br>TTCCTTTTTGTTGCGCATTTCTATATACGCTTCTGCAACATCTGAGAAATGTGCCATGATTTGCC<br>TAAGTGTTGGTTTTGCATTCTCAACGATTGGTTTCAGTGGGTATTCGACTTGTTCATCTCCATCC<br>ATCATAACCCAAACTC |
| RNA RC Long | AACUUCAUAAAAGUCAAAAGCAUAGCGAGCCAAACUUCCAUCGCGCAGAUUACGAACUAAA<br>CCAUAUCGUGGCAUAUAUGGUUCCUUUUUGUUGCGCAUUUCUAUAUACGCUUCUGCAACAU<br>CUGAGAAAUGUGCCAUGAUUUGCCUAAGUGUUGGUUUUGCAUUCUCAACGAUUGGUUUCA<br>GUGGGUAUUCGACUUGUUCAUCUCCAUCCAUCAUAACCCAAACUC |

**Supplementary Table S6: DNA and RNA sequences used for screening PVY 5 to 8.** The bold sequence represents the extended T7 promoter used for the generation of each RNA via *in vitro* transcription.

| Place | Method | Cost in local currency | Cost in US dollars | % cost reduction from the value above | to UK price | to Chile price |
|---|---|---|---|---|---|---|
| Chile, PUC | PURExpress | 5016 CLP | 7.76 | | 1.2 | NA |
| | 3-PGA | 56 CLP | 0.086 | 98.9 | 1.93 | NA |
| | MDX | 45 CLP | 0.069 | 20 | 2 | NA |
| UK, UniCam | PURExpress | 4.41 GBP | 6.5 | | NA | 0.83 |
| | 3-PGA | 0.016 GBP | 0.045 | 99 | NA | 0.52 |
| | MDX | 0.012 GBP | 0.036 | 23 | NA | 0.5 |

**Supplementary Table S7: Cost breakdown analysis of cell-free reactions performed in Chile and UK (Cambridge University).** All the reagents needed for the production of maltodextrin (MDX) or 3-PGA methods, along with the reagents needed for extract production were used to calculate a cost per reaction (considering 5 ul reaction volume). Exchange rates as per in the $8^{th}$ of June 2020 were used to convert local currency to USD. This value was used to compare cost reductions with respect to the commercial alternative PURExpress (also considering 5 ul reaction volume) as well as for cost differences between places and methods.

| Item | Supplier | Cat number | Quantity (g) | cost (GBP) | % on cost per reaction |
|---|---|---|---|---|---|
| COENZYME A HYDRATE | Sigma/Merck | C4282-100MG | 0.1 | 220.8 | 7.204 |
| B-NICOTINAMIDE ADENINE DINUCLEOTIDE HYDR | Sigma/Merck | N6522-250MG | 0.25 | 33.2 | 0.49 |
| L-AMINO ACIDS KIT | Sigma/Merck | LAA21-1KT | 20 | 300 | 2.301 |
| L-Arginine | Sigma/Merck | 11009-25G-F | 25 | 25.82 | 0.01 |
| L-Cysteine | Sigma/Merck | 30089-25G | 25 | 43.02 | 0.012 |
| L-Histidine | Sigma/Merck | 53319-25G | 25 | 39.69 | 0.014 |
| L-GLUTAMIC ACID HEMIMAGNESIUM SALT TETRA | Sigma/Merck | 49605-250G | 250 | 32.81 | 0.002 |
| L-GLUTAMIC ACID POTASSIUM SALT MONO& | Sigma/Merck | G1149-100G | 100 | 33.15 | 0.062 |
| FOLINIC ACID CALCIUM | Sigma/Merck | F7878-100MG | 0.1 | 63.75 | 0.363 |
| GTP, DISODIUM SALT | Roche | 10106399001 | 0.25 | 102.85 | 5.278 |
| ADENOSINE 5'-TRIPHOSPHATE DIPOTASSIUM SALT | Sigma/Merck | A8937-1G | 1 | 42.3 | 0.605 |
| CYTIDINE 5'-TRIPHOSPHATE DISODIUM SALT | Sigma/Merck | C1506-100MG | 0.1 | 84.15 | 10.153 |
| ADENOSINE 3":5"-CYCLIC MONOPHOSPHATEFR | Sigma/Merck | A9501-1G | 1 | 146.2 | 0.595 |
| TRNA, FROM E.COLI MRE 600, 100 MG | Roche | 10109541001 | 0.1 | 119.85 | 0.392 |
| 3-PGA | Sigma/Merck | P8877 | 1 | 205.33 | 23.165 |

| | | | | | |
|---|---|---|---|---|---|
| URIDINE 5'-TRIPHOSPHATE TRISODIUM SALT D | Sigma/Merck | 94370-250MG | 0.25 | 43.43 | 2.331 |
| SPERMIDINE, FOR MOLECULAR BIOLOGY | Sigma/Merck | 85558-5G | 5 | 126 | 0.494 |
| HEPES | Sigma/Merck | H6147-25G | 25 | 53.47 | 0.417 |
| PEG 8000 | Promega | V3011 | 500 | 53.25 | 0 |
| Crude Extract | In-house (see Supplementary Table S9) | NA | NA | 466 | 46.113 |

**Supplementary Table S8: Cost breakdown analysis of cell-free reactions produced in UK using 3-PGA as energy source.** 3-PGA reagent is shown highlighted in yellow.

| Item | Supplier | Cat number | Quantity (g) | cost (GBP) | % on cost per reaction |
|---|---|---|---|---|---|
| COENZYME A HYDRATE | Sigma/Merck | C4282-100MG | 0.1 | 220.8 | 9.367 |
| B-NICOTINAMIDE ADENINE DINUCLEOTIDE HYDRATE | Sigma/Merck | N6522-250MG | 0.25 | 33.2 | 0.637 |
| L-AMINO ACIDS KIT | Sigma/Merck | LAA21-1KT | 20 | 300 | 2.992 |
| L-Arginine | Sigma/Merck | 11009-25G-F | 25 | 25.82 | 0.013 |
| L-Cysteine | Sigma/Merck | 30089-25G | 25 | 43.02 | 0.015 |
| L-Histidine | Sigma/Merck | 53319-25G | 25 | 39.69 | 0.018 |
| L-GLUTAMIC ACID HEMIMAGNESIUM SALT | Sigma/Merck | 49605-250G | 250 | 32.81 | 0.003 |
| L-GLUTAMIC ACID POTASSIUM SALT | Sigma/Merck | G1149-100G | 100 | 33.15 | 0.08 |
| FOLINIC ACID CALCIUM | Sigma/Merck | F7878-100MG | 0.1 | 63.75 | 0.471 |
| GTP, DISODIUM SALT | Roche | 10106399001 | 0.25 | 102.85 | 6.863 |
| ADENOSINE 5'-TRIPHOSPHATE DIPOTASSIUM SALT | Sigma/Merck | A8937-1G | 1 | 42.3 | 0.787 |
| CYTIDINE 5'-TRIPHOSPHATE DISODIUM SALT | Sigma/Merck | C1506-100MG | 0.1 | 84.15 | 13.202 |
| ADENOSINE 3'':5''-CYCLIC MONOPHOSPHATEFR | Sigma/Merck | A9501-1G | 1 | 146.2 | 0.774 |
| TRNA, FROM E.COLI MRE 600, 100 MG | Roche | 10109541001 | 0.1 | 119.85 | 0.51 |

| | | | | | |
|---|---|---|---|---|---|
| SODIUM POLYPHOSPHATE, CRYSTALS (HMP) | Sigma/Merck | 305553-25G | 25 | 29.33 | 0.015 |
| MALTODEXTRIN, DEXTROSE EQUIVALENT 4.0-7. | Sigma/Merck | 419672-100G | 100 | 30.24 | 0.078 |
| URIDINE 5'-TRIPHOSPHATE TRISODIUM SALT D | Sigma/Merck | 94370-250MG | 0.25 | 43.43 | 3.03 |
| SPERMIDINE, FOR MOLECULAR BIOLOGY | Sigma/Merck | 85558-5G | 5 | 126 | 0.643 |
| HEPES | Sigma/Merck | H6147-25G | 25 | 53.47 | 0.542 |
| PEG 8000 | Promega | V3011 | 500 | 53.25 | 0 |
| Crude Extract | In-house (see Supplementary Table S9) | NA | NA | 466.1 | 59.961 |

**Supplementary Table S9: Cost breakdown analysis of cell-free reactions produced in UK using maltodextrin and polyphosphates as energy source**. Reagents specific to the maltodextrin energy solution are highlighted in yellow.

| Item | Supplier | Cat number | Quantity (g) | cost (GBP) | % on cost per reaction |
|---|---|---|---|---|---|
| 2xYTG medium (YT powder) | Fisher Scientific Ltd/MP Biomedicals | 11357669 | 454 | 50.16 | 14.57 |
| 2xYTG medium (Potassium Phosphate Dibasic 1M - K2HPO4) | Sigma | P8584-1L | 1L | 34.8 | 5.92 |
| 2xYTG medium (Potassium Phosphate Monobasic 1M - KH2PO4) | Sigma | P8709-1L | 1L | 33.92 | 3.17 |
| 2xYTG medium (D-glucose ) | Sigma | G8270-1KG | 1000 | 27.14 | 2.08 |
| Induction (IPTG) | Thermo Scientific | 15763552 | 1 | 34.22 | 34.69 |
| Cell extract separation (Micro Bio-Spin Chromatography Columns, empty, 100) | BioRad | 7326204 | 100 | 97.11 | 30.25 |
| S30B Buffer (DITHIOTHREITOL molecular grade) | Sigma | D9779-5G | 5 | 90.96 | 5.37 |
| S30B Buffer (L-GLUTAMIC ACID HEMIMAGNESIUM SALT TETRA) | Sigma/Merck | 49605-250G | 250 | 32.81 | 0.1 |
| S30B Buffer (L-GLUTAMIC ACID POTASSIUM SALT MONO) | Sigma/Merck | G1149-100G | 100 | 33.15 | 3.81 |
| S30B Buffer (TRIS) | Fisher Scientific | BP1521 | 1000 | 31.83 | 0.04 |

**Supplementary Table S10: Cost breakdown analysis of cell extract preparation including culture, induction, and lysis as costs listing in U. of Cambridge (UK).**

| Item | Supplier | Distributor in Chile | Cat number | Quantity (g) | initial cost (CLP) | % on cost per reaction |
|---|---|---|---|---|---|---|
| COENZYME A HYDRATE | Sigma/Merck | Sigma-Aldrich Ltda | C4282-35MG | 0.035 | 194,000 | 9.396 |
| B-NICOTINAMIDE ADENINE DINUCLEOTIDE HYDR | Sigma/Merck | GenExpress | N6522-1G | 1 | 27,904 | 0.053 |
| L-AMINO ACIDS KIT | Sigma/Merck | Sigma-Aldrich Ltda | LAA21-1KT | 20 | 491,000 | 1.957 |
| L-Cysteine | Sigma/Merck | Sigma-Aldrich Ltda | 1028380100 | 100 | 57,206 | 0.002 |
| L-Histidine | Sigma/Merck | Sigma-Aldrich Ltda | 1043510100 | 100 | 103,447 | 0.005 |
| L-GLUTAMIC ACID HEMIMAGNESIUM SALT | Sigma/Merck | Sigma-Aldrich Ltda | 49605-250G | 250 | 74,000 | 0.003 |
| L-GLUTAMIC ACID POTASSIUM SALT MONO | Sigma/Merck | Sigma-Aldrich Ltda | G1149-100G | 100 | 74,000 | 0.072 |
| FOLINIC ACID CALCIUM | Sigma/Merck | Fermelo | F7878-100MG | 0.1 | 71,000 | 0.21 |
| GTP, DISODIUM SALT | Sigma/Merck | Sigma-Aldrich Ltda | A2383-250MG | 0.25 | 136,000 | 3.626 |
| ADENOSINE 5'-TRIPHOSPHATE DIPOTASSIUM SALT | Sigma/Merck | Sigma-Aldrich Ltda | A8937-1G | 1 | 80,000 | 5.946 |
| CYTIDINE 5'-TRIPHOSPHATE DISODIUM SALT | Affymetrix/USB | Biosonda | 14121-100MG | 0.1 | 195,468 | 12.253 |
| ADENOSINE 3":5"-CYCLIC MONOPHOSPHATEFR | Sigma/Merck | Fermelo | A9501-100MG | 0.1 | 108,000 | 2.283 |
| TRNA, FROM E.COLI MRE 600, 100 MG | Roche | Sigma-Aldrich Ltda | MRE600 | 0.1 | 143,000 | 0.243 |

| | | | | | | |
|---|---|---|---|---|---|---|
| 3-PGA | Sigma/Merck | Sigma-Aldrich Ltda | P8877 | 1 | 394,000 | 23.095 |
| URIDINE 5'-TRIPHOSPHATE TRISODIUM SALT D | Affymetrix/USB | GenExpress | 23160-100MG | 0.1 | 75,000 | 5.228 |
| SPERMIDINE, FOR MOLECULAR BIOLOGY | Sigma/Merck | Sigma-Aldrich Ltda | 85558-1G | 1 | 81,000 | 0.826 |
| HEPES | Sigma/Merck | Winkler | H6147-100G | 100 | 61,250 | 0.062 |
| PEG 8000 | Promega | Galenica | V30111 | 500 | 59,000 | 0 |
| Crude extract | In-house (see Supplementary Table S12) | NA | NA | NA | 1,047,500 | 34.74 |

**Supplementary Table S11: Cost breakdown analysis of cell-free reactions produced in Chile using 3-PGA as energy source.** 3-PGA is highlighted in yellow.

| Item | Supplier | Distributor in Chile | Cat number | Quantity (g) | initial cost (CLP) | % cost per reaction |
|---|---|---|---|---|---|---|
| COENZYME A HYDRATE | Sigma/Merck | Sigma-Aldrich Ltda | C4282-35MG | 0.035 | 194,000 | 12.396 |
| B-NICOTINAMIDE ADENINE DINUCLEOTIDE HYDR | Sigma/Merck | GenExpress | N6522-1G | 1 | 27,904 | 0.071 |
| L-AMINO ACIDS KIT | Sigma/Merck | Sigma-Aldrich Ltda | LAA21-1KT | 20 | 491,000 | 2.582 |
| L-Cysteine | Sigma/Merck | Sigma-Aldrich Ltda | 1028380100 | 100 | 57,206 | 0.003 |
| L-Histidine | Sigma/Merck | Sigma-Aldrich Ltda | 1043510100 | 100 | 103,447 | 0.006 |
| L-GLUTAMIC ACID HEMIMAGNESIUM SALT | Santa Cruz | Fermelo | SC228394 | 150 | 47,500 | 0.002 |
| L-GLUTAMIC ACID POTASSIUM SALT MONO | Santa Cruz | Fermelo | SC250217 | 100 | 39,500 | 0.05 |
| FOLINIC ACID CALCIUM | Sigma/Merck | Fermelo | F7878-100MG | 0.1 | 71,000 | 0.277 |
| GTP, DISODIUM SALT | Sigma/Merck | Sigma-Aldrich Ltda | A2383-250MG | 0.25 | 136,000 | 4.784 |
| ADENOSINE 5'-TRIPHOSPHATE DIPOTASSIUM SALT | Sigma/Merck | Sigma-Aldrich Ltda | A8937-1G | 1 | 80,000 | 7.845 |
| CYTIDINE 5'-TRIPHOSPHATE DISODIUM SALT | Affymetrix/USB | Biosonda | 14121-100MG | 0.1 | 195,468 | 16.165 |
| ADENOSINE 3":5"-CYCLIC MONOPHOSPHATEFR | Sigma/Merck | Fermelo | A9501-100MG | 0.1 | 108,000 | 3.012 |

| | | | | | | |
|---|---|---|---|---|---|---|
| TRNA, FROM E.COLI MRE 600, 100 MG | Roche | Sigma-Aldrich Ltda | MRE600 | 0.1 | 143,000 | 0.321 |
| SODIUM POLYPHOSPHATE, CRYSTALS (HMP) | Sigma/Merck | Sigma-Aldrich Ltda | 305553-25G | 25 | 30,553 | 0.008 |
| MALTODEXTRIN, DEXTROSE EQUIVALENT 4.0-7. | Sigma/Merck | Sigma-Aldrich Ltda | 419672-100G | 100 | 55,000 | 0.075 |
| URIDINE 5'-TRIPHOSPHATE TRISODIUM SALT D | Affymetrix/USB | GenExpress | 23160-100MG | 0.1 | 58,733 | 5.401 |
| SPERMIDINE, FOR MOLECULAR BIOLOGY | Sigma/Merck | Sigma-Aldrich Ltda | 85558-1G | 1 | 81,000 | 1.089 |
| HEPES | Sigma/Merck | Winkler | H6147-100G | 100 | 61,250 | 0.082 |
| PEG 8000 | Promega | Galenica | V30111 | 500 | 59,000 | 0 |
| Crude extract | In-house (see Supplementary Table S12) | NA | NA | NA | 1,047,500 | 45.832 |

**Supplementary Table S12: Cost breakdown analysis of cell-free reactions produced in Chile using maltodextrin and polyphosphates as energy source.** Reagents needed for maltodextrin energy source are highlighted in yellow.

| Item | Supplier | Distributor in Chile | Cat number | Quantity (g) | initial cost (CLP) | % cost per rxn |
|---|---|---|---|---|---|---|
| 2xYTG medium (YT powder) | MP Miomedicals | Galenica | 113012022 | 454 | 56,000 | 11.22 |
| 2xYTG medium (Potassium Phosphate Dibasic 1M - K2HPO4) | Sigma | Sigma-Aldrich | P8584-1L | 1L | 36,700 | 4.31 |
| 2xYTG medium (Potassium Phosphate Monobasic 1M - KH2PO4) | Sigma | Sigma-Aldrich | P8709-1L | 1L | 42,400 | 2.74 |
| 2xYTG medium (D-glucose ) | Sigma | Sigma-Aldrich | G5767 | 500 | 50,900 | 5.38 |
| Induction (IPTG) | Promega | Fermelo | V3951 | 5 | 199,500 | 27.89 |
| Cell extract separation (Micro Bio-Spin Chromatography Columns, empty) | Bio Rad | Galenica | 7326204 | 100 | 198,000 | 42.54 |
| S30B Buffer (DITHIOTHREITOL molecular grade) | Promega | Fermelo | V3155 | 25 | 310,000 | 2.53 |
| S30B Buffer (L-GLUTAMIC ACID HEMIMAGNESIUM SALT TETRA) | Santa Cruz | Fermelo | SC228394 | 150 | 47,500 | 0.16 |
| S30B Buffer (L-GLUTAMIC ACID POTASSIUM SALT MONO) | Santa Cruz | Fermelo | SC250217 | 100 | 39,500 | 3.13 |
| S30B Buffer (TRIS) | Bio Rad | Galenica | BP 1 52 1 | 500 | 67,000 | 0.11 |

**Supplementary Table S13: Cost breakdown analysis of cell extract preparation including culture, induction and lysis as prices listing in PUC university in Chile.**
